# Affiliative behaviours regulate allostasis development and shape biobehavioural trajectories in horse

**DOI:** 10.1101/2025.01.14.632964

**Authors:** Mathilde Valenchon, Fabrice Reigner, Gaëlle Lefort, Hans Adriaensen, Amandine Gesbert, Philippe Barrière, Yvan Gaude, Frederic Elleboudt, Isabelle Lévy, Camille Ducluzeau, Joëlle Dupont, Anne-Lyse Lainé, Ivy Uszynski, Hugues Dardente, Cyril Poupon, Léa Lansade, Ludovic Calandreau, Matthieu Keller, David André Barrière

## Abstract

Social interactions shape both physiological and behavioural development of offspring and poor care/early caregiver loss are known to promote negative outcomes in adulthood in both animals and humans. How affiliative behaviours impact future development of offspring remains unknown. Here, we used *Equus caballus* (domestic horse) as a model to investigate this question. By coupling magnetic resonance imaging, longitudinal biobehavioural assessment and advanced multivariate statistical modelling we found that maternal presence during childhood promotes maturation of brain territories involved in both social behaviour (anterior cingulate, retrosplenial cortex) and physiological regulation (hypothalamus, amygdala). Additionally, we found that offsprings benefiting from prolonged maternal presence showed higher default mode network (DMN) connectivity, improved social competences, more efficient feeding behaviours, and metabolic profiles. The present study underscores the salient role of social interactions for the development of allostatic regulation in offspring.

## Main

In large social mammals including humans, caregivers-typically parents-provide care through a species-specific immature period beyond the lactation period, to ensure the acquisition of social skills necessary for survival and reproductive fitness of the offspring^1–3^. Caregiver death/loss or poor caregiving during this socially immature period, are known to promote negative physiological and behavioural outcomes in adulthood in both animals and humans^2,4^. In wild primates considered as altricial (ex. Mountain gorilla), longitudinal studies have shown that offspring is more likely to die early in adulthood when mothers experienced poor physical conditions, and display poorer abilities to raise their progeny^1,2^. In precocial species such as horse, previous studies have shown that an abrupt rupture of the maternal bound around 4-6 months of age induces long-lasting emotional/social outcomes, oral/locomotor stereotypies and physiological impairments of cortisol/lipid metabolism associated with reduction of telomer lengths in offspring^5–7^. Hence, regardless of brain maturity at birth, its seems that in large social mammals, affiliative behaviours provide social interactions salient enough to foster the development of regulatory systems vital to control physiology and behaviours in adulthood^3,4,8^.

During the last decade, allostasis, a brain-centred concept, has been invoked to explain this phenomenon^4,9–13^. Allostasis posits that brain is dedicated to make predictive and anticipatory regulations of the internal milieu through continuous integration of internal/external cues necessary for survival^9,14,15^. In adult human brain, allostasis would be supported by a complex system integrating cortices (cingulate, insula, prefrontal), hippocampus, striatum, hypothalamus, amygdala, periaqueductal grey and parabrachial nucleus^16–19^. In this system, cortical areas are embedded within domain-general core networks such as the “default mode network” (DMN), in which intero/exteroceptive inputs are constantly integrated to build predictive models. These models would then be compared by primary sensory cortices, and the difference (i.e. error prediction) used as a final output to tune the autonomic system. This allostatic regulatory system would control the activity of peripheral organs (e.g. adrenals, etc.) through the release of endocrine hormones by hypothalamus, but also brainstem regions (e.g. PAG), which regulate homeostasis (e.g. heart rate) and behaviours (e.g. fight/flight). Importantly, the level of error prediction, encoded by striatal dopamine levels, would support learning mechanisms used to refine forthcoming predictions and minimize energy requirement for future adaptations^4,20^. Therefore, the allostatic regulation system is an acquired system, developed during infancy and further refined during lifetime through physical and social experiences. This system is key for daily physiological/behavioural adaptations of an individual to navigate in his environnement^3,4,8^.

To date, literature supports the hypothesis that the initial caregiver-offspring dyad is a critical scheme in which affiliative behaviours, fuelled by the maternal bond, provide social cues necessary for the maturation of the allostatic regulatory system during the socially immature period^1–3^. This process would be shared by large social mammals as being vital for adaptation of offspring to a challenging physical/social environment. However, we still don’t know how social interactions engage these brain-maturing processes, what these processes and their neural substrates are, and how their eventually shape physiological and behavioural trajectories during development.

### *Equus caballus* to study brain development and biobehavioural trajectories

We chose to evaluate in *Equus caballus* (domestic horse) how affiliative behaviours during infancy influence the development of the allostasis regulating system and further shape biobehavioural trajectories of offspring. The mother-offspring dyad ontogeny in horse shares many features with large social mammals’ species such as ungulates, cetaceans and primates including humans, which makes it an excellent model for such investigation. Horses typically have one/two offspring(s) per gestation, provide an extended period of parental care (>1 year), and mothers rapidly establish a strong, long-lasting and individualized affiliative bond with their offspring^21^. To test our hypothesis, we developed a holistic approach based on *in vivo* magnetic resonance imaging (MRI) to study brain structure and function in horse offspring associated with a longitudinal collection of behavioural and physiological data between 6 to 13 months of age (**Fig.1a**). In horse ontogeny, this timeframe is considered has a “mid-childhood” period starting when offsprings are able to feed themselves and control their physiology (around 6 months) but are still socially dependent of their mothers, and terminating with the onset of sexual maturation, which marks the beginning of the adolescence period (around 13 months)^22^. During this timeframe, we applied our multimodal approach to two animal groups (total of 24 animals) herded in the same conditions (indoor collective stalls with free access to an outdoor paddock). The first group named “*maternal presence*” was composed of unweaned offsprings (6 males/6 females), which have been raised with their mothers (12 mares). The second group named “*maternal absence*” was composed of 12 weaned offsprings (6 males/6 females), which have been separated from their mothers at the beginning of the experiment at 6 months of age (**Fig.1b**). In total, 36 animals (offsprings plus mothers) were herded together. This procedure allow to study the impact of caregiver loss on brain development and biobehavioural trajectories until adolescence and minimize biases induced by sexual hormones but also stress and/or emotional contagion, through the promotion of a socially enriched environment for weaned animals. To explore the impact on the foals’ brain development, multimodal MRI data (functional, diffusion and anatomical) were collected 1 month after weaning procedure in both groups to evaluate both functional and microstructural variations between groups (**Fig.1c**). *In vivo* imaging data have been used to create the first *in vivo* brain template and atlas of *equidae*, the <u>Turone</u> <u>Equine Brain Template and Atlas</u>, which was used for MRI data analysis (see *supplementary material* **Fig. S1** and **Table S1**). To explore the role of extended maternal care onto the foals’ behavioural development, multiple evaluations have been performed 1-, 3-, and 7-months after weaning procedure, to assess activity budgets, spatial proximity, centrality and spontaneous behavioural activity of each animal. Additionally, sociability and fearfulness were assessed through standardised behavioural tests 7-months post-treatment (**Fig.1d**). Finally, to ascertain the impact of the maternal presence on the offsprings’ physiological development, animals were weighed and blood sampled at 1-, 3- and 7-months to assess plasma levels of basal cortisol, oxytocin, glycemia, cholesterol, IGF-1, and insulin (**Fig.1e**).

**Fig1.**
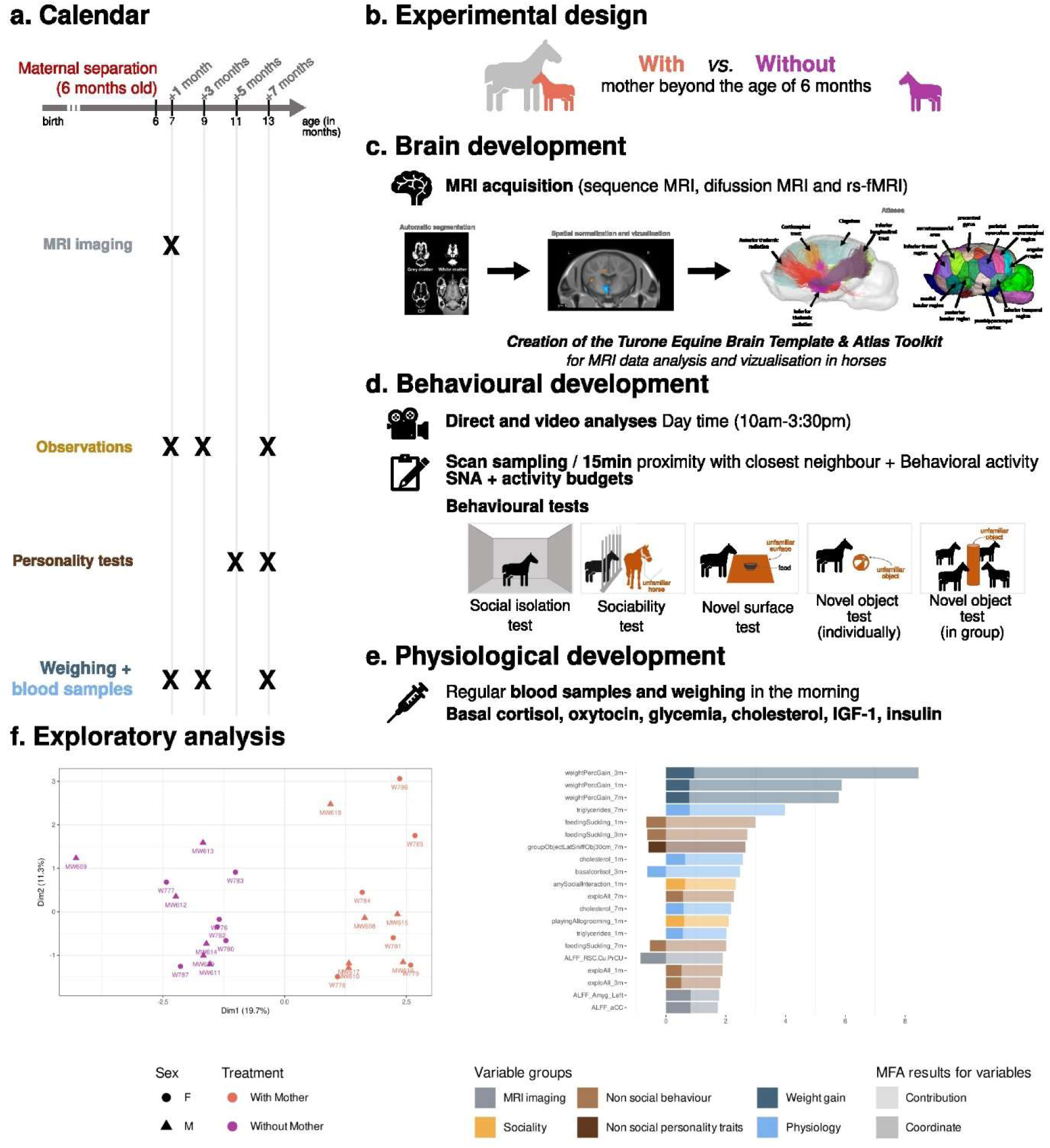
Experimental design of our comprehensive and longitudinal study to assess how a maternal presence beyond 6 months of age influences brain, behavioural, and physiological development. (**a**) Timeline and (**b**) overview of the experimental protocol, with (**c-e**) representing the main categories of the recorded parameters over time including (**c**) brain development (functional/microstructural structural MRI brain metrics), (**d**) behavioural development (behavioural observations and tests), and (**e**) physiological development (hormonal and metabolic profiling). A comprehensive visualisation of the dataset is shown in (**f-g**) using an integrative Multiple Factor Analysis where (**f**) shows the distribution of individuals along the first two axes of the MFA, and (**g**) displays the variables that contribute the most to the first MFA axis, which discriminates between the “with mother” and “without mother” animals.

### Multimodal approach discriminates the effect of prolonged maternal presence

At the end of the experiment, a total of 76 variables were collected for 23 animals as one female belonging to the “with mother” group leave the experiment for medical reasons. From this dataset, we first conducted a holistic exploratory analysis to determine the potential for discriminating individuals based on treatment (“*maternal presence*” versus “*maternal absence*”). To achieve this, we performed a Multiple Factor Analysis (MFA) on all parameters for which variables were grouped into equally weighed sets for analysis, and six groups of variables were defined: *weight gain, non-social behaviour, non-social personality traits, sociality, MRI imaging*, and *physiology*. MFA analysis showed a clear separation between the “*maternal presence*” (orange) and “*maternal absence*” (purple) groups across the first dimension, demonstrating the multidimensional impact of extended maternal care onto offsprings development (**Fig. 1f**). Physiological variables exhibit the most significant disparities between groups, followed by behavioural variables and, finally, specific brain metrics from brain regions involved in the allostasis regulatory system (**Fig. 1g**).

### Prolonged maternal presence shapes brain microstructural organization

Using our <u>TEBTA resources</u> (**Fig. 2**) we investigated the effects of maternal presence on brain microstructural organisation of the offspring. We first evaluated the difference in grey matter concentration (GMC) using a voxel-based morphometry approach (VBM). VBM is a thoroughly validated technique of imaging analysis that provides an unbiased and comprehensive assessment of anatomical differences throughout the brain grey matter in both animal and human brains^23–26^. Using MRI T_1w_ anatomical acquisitions, we estimated the GMC maps, which provided a global estimation of the GMC of the brain for each animal. Comparison of GMC maps revealed significantly lower GMC values (Student t-test, voxel-level threshold P < 0.05, t_(21)_ = 1.7207, FDR-corrected) in animals belonging to the “*maternal presence*” group for the amygdala (amyg), the hippocampus (hippo), and the medial cingulate cortex (mCC), and substantially higher values in both the thalamus (thal) and the hypothalamus (hypo) (**Fig. 3**). Although VBM is a highly sensitive method to study brain microstructural modifications, its specificity is limited and functional significance of such microstructural modifications still is a matter of debate^27^. Therefore, to better characterise these microstructural modifications, we used a multi-shell 3D diffusion MRI approach (dMRI) optimised for neurite orientation dispersion and density imaging (NODDI), according to the model of Zhang *et al*^28^. From this imaging dataset, we extracted the fractional anisotropy (FA), along with three scalar parameters from the NODDI model: the intra-neurite fraction (inf), also known as the neurite density index representing axons and possibly dendritic processes, the orientation dispersion index (ODI, where 0 indicates perfectly aligned straight axons, and 1 indicates fully isotropic axons), and finally, the isotropic fraction (isoFrac, the extra-neurite compartment which encompasses everything except neurites and free water, such as microglia, astrocytes, oligodendrocytes, neuronal somas, ependymal cells, extracellular matrices, and vascular structures)^29^. Comparison of FA maps showed significantly lower FA values (Student t-test, voxel-level threshold p < 0.05, t t_(21)_ = 1.7207, FDR-corrected) within the anterior cingulate cortex (aCC), the cuneus/precuneus (Cu/PrCu); the prefrontal cortex (pFC); and the retrosplenial cortex (RSC) for animals belonging to the “*maternal presence*” group (**Fig. 4a**). Strikingly, significantly higher ODI values (Student t-test, voxel-level threshold p < 0.05, t_(21)_ = 1.7207, FDR-corrected) were observed in similar regions for animals belonging to the “*maternal presence*” (**Fig. 4b**). Finally, significantly lower isoFrac values (Student t-test, voxel-level threshold p < 0.05, t_(21)_ = 1.7207, FDR-corrected) were found within the cuneus/precuneus (Cu/PrCu), the prefrontal cortex (pFC) and hypothalamus (hypo) for animals belonging to the “*maternal presence*” group (**Fig. 4c**). Altogether, both VBM and dMRI revealed that offsprings benefiting from prolonged maternal presence, exhibited significant microstructural modifications within pivotal brain structures mainly involved in physiological (thalamus) and behavioural (aCC/mCC, Cu/PrCu, pFC and RSC) regulations or both (hypothalamus).

**Fig 2.**
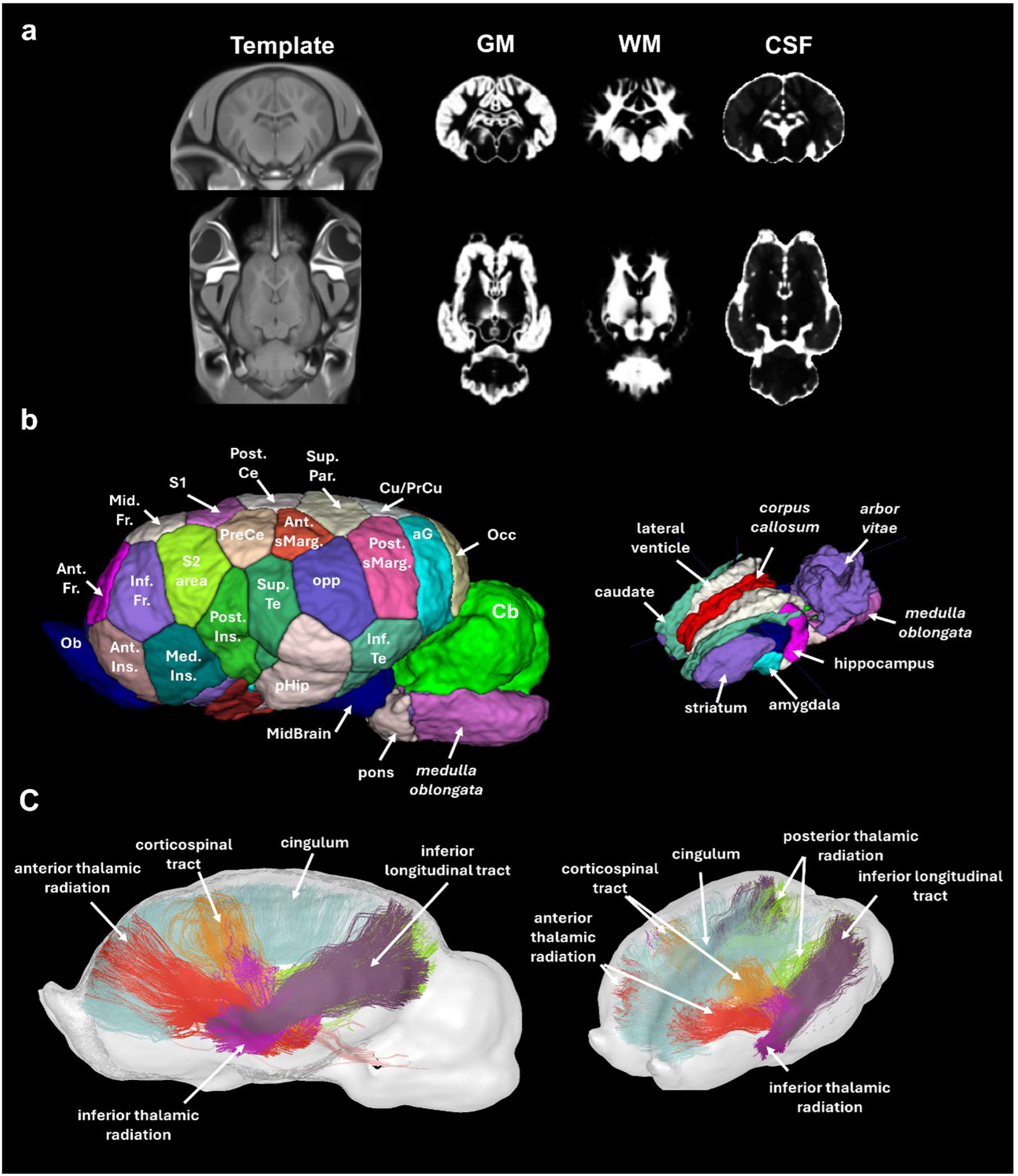
The Turone Equine Brain Template & Atlas. (**a**) Sagittal and axial brain slices of Turone equine template and its associated tissue probability maps of gray matter (GM), white matter (WM) and cerebrospinal fluid (CSF). (**b**) Lateral and top views of the Turone equine brain atlas show the results of the multi-animal dictionary learning statistical approach for cortical parcellation using functional imaging data (left) and the results of the manual segmentation of the subcortical regions (right). (**c**) Lateral and top views of the Turone equine brain fibre atlas show the results of the manual fibre clustering using diffusion imaging data. The complete list of the region of interest and their full names are freely available with the TEBTA resource on <u>Zenodo website</u>.

**Fig 3.**
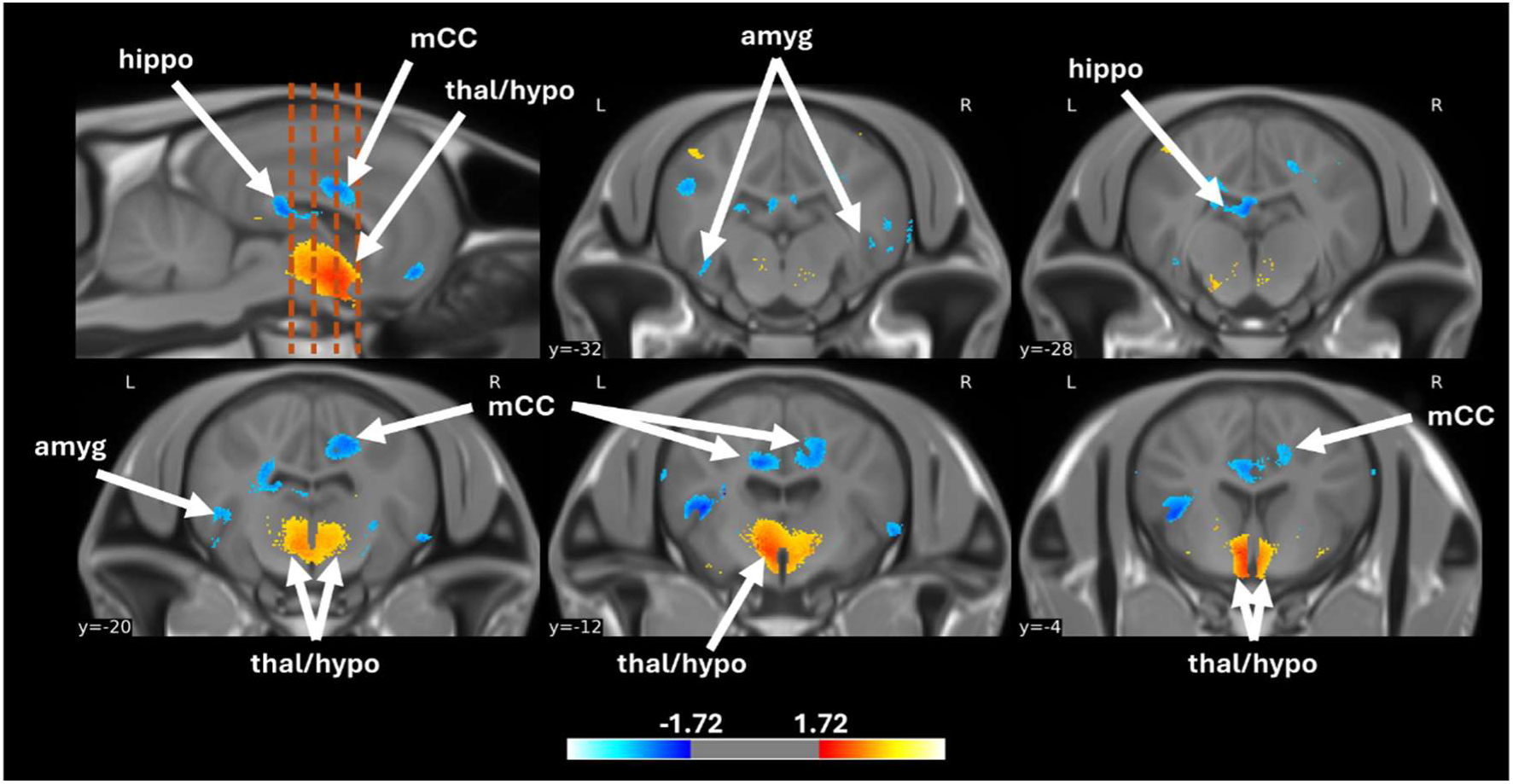
Effect of prolonged maternal presence on grey matter concentration in the equine brain (*with mother > without mother*). Sagittal and axial brain slices showing grey matter concentration (GMC) differences between groups. Orange dashed lines represent the approximate positions of axials brain slices. Data obtained from SPM unpaired Student t-test analysis using a voxel-level threshold p < 0.05, t (21) = 1.7207, FDR-corrected. *Abbreviations: amyg = amygdala; hippo = hippocampus; hypo = hypothalamus; mCC = medial cingulate cortex, thal = thalamus*.

**Fig 4.**
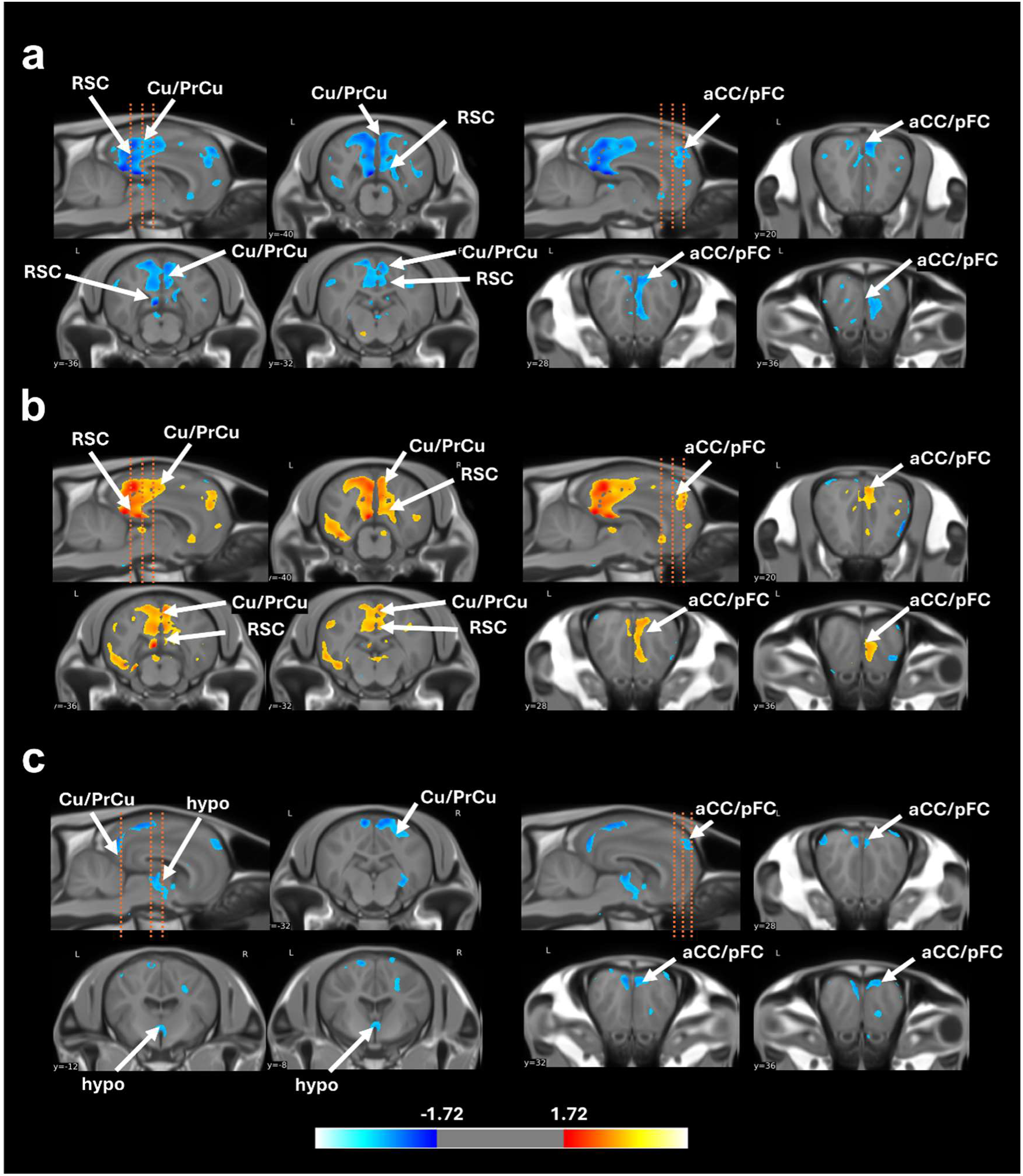
Effect of a maternal prolonged maternal presence on the cytoarchitecture of the equine brain (*with mother > without mother*). Sagittal and axial brain slices showing modification of (**a**) fractional anisotropy (FA), (**b**) orientation dispersion index (ODI) and (**c**) isotropic fraction between groups. Orange dashed lines represent the approximate positions of axial brain slices. Data obtained from SPM unpaired Student t-test analysis using a voxel-level threshold p < 0.05, t (21) = 1.7207, FDR-corrected. *Abbreviations: aCC = anterior cingulate cortex; Cu/PrCu = cuneus/precuneus; hypo = hypothalamus; pFC = prefrontal cortex; RSC = retrosplenial cortex*.

### Prolonged maternal presence shapes brain functioning

To study the impact of such microstructural modifications on brain functioning, we used functional MRI (fMRI) optimised for resting-state analysis. From these data, we computed the fractional amplitude of low-frequency fluctuation maps (fALFF, 0.01–0.08 Hz), a highly stable, specific and sensitive index of resting-state fMRI used to characterise spontaneous brain activity^30,31^. Firstly, comparison of fALFF revealed significant variations (Student t-test, voxel-level threshold p < 0.05, t_(21)_ = 1.7207, FDR-corrected) in spontaneous neuronal activity within previously highlighted regions. Specifically, significant lower fALFF values were observed within cuneus/precuneus (Cu/PrCu), retrosplenial cortex (RSC), prefrontal cortex (pFC) of animals belonging to the “*maternal presence*” group. Moreover, significant higher fALFF values were observed within the anterior cingulate cortex (aCC), the periaqueductal grey substance (PAG), and the ventral hippocampus (vHipp) in the brains of animals belonging to the “*maternal presence*” group (**Fig. 5a-b**).

**Fig 5.**
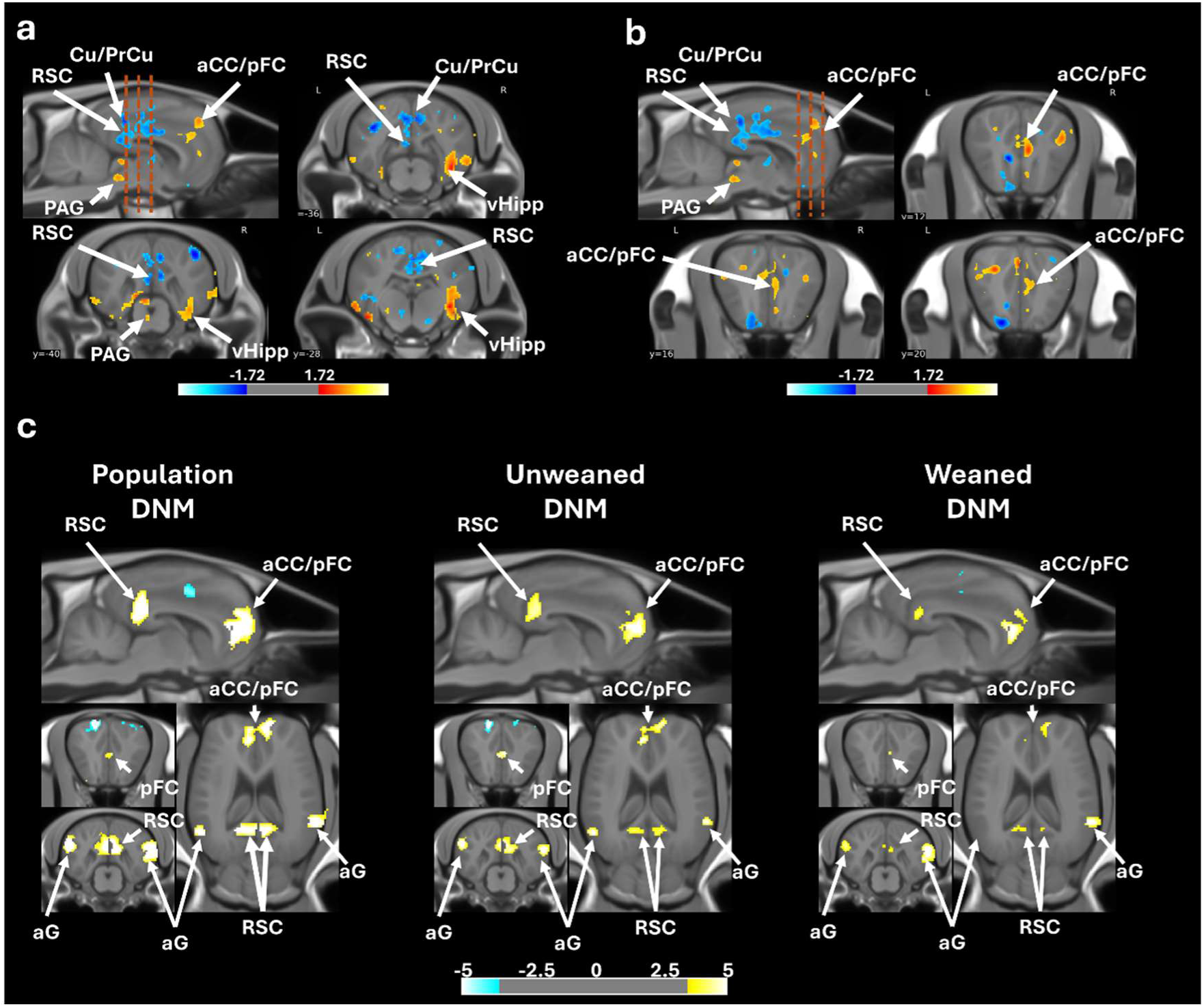
Effect of a maternal prolonged maternal presence on functional organisation of the equine brain (*with mother > without mother*). Sagittal and axial brain slices showing modifications of the amplitude of low frequency fluctuations (ALFF) in posterior (a) and anterior (b) between groups. Orange dashed lines represent the approximate positions of axials brain slices. The default mode network (DMN) is identified in the domestic horse by calculating the mean Pearson correlation coefficient between time-series of spontaneous variations of BOLD (Blood Oxygen Level Dependent) signal, extracted from anatomically defined regions of interest (ROIs) which are functionally homologous to regions involved in DMN in numerous species (aG, aCC/pFC and RSC) and in which spontaneous variations of the BOLD signal is highly correlated at rest between them. For (**a**) and (**b**) figures, orange dashed lines represent the approximate positions of axials brain slices. Data obtained from SPM unpaired Student t-test analysis using a voxel-level threshold p < 0.05, t (21) = 1.7207, FDR-corrected. For (**c**) data were obtained from SPM multiple regression analysis using a voxel-level threshold p < 0.001, t (21) = 3.5272, FDR-corrected. *Abbreviations: aG = angular gyrus; aCC = anterior cingulate cortex; Cu/PrCu = cuneus/precuneus; PAG = periaqueductal grey; vHipp = ventral hippocampus; pFC = prefrontal cortex; RSC = retrosplenial cortex*.

This overlap between both functional and microstructural modifications within the Cu/PrCu, the RSC, the pFC and the aCC - which are known to be key players in the default mode network (DMN), a large-scale brain network involved in socio-cognitive development in humans - suggests that prolonged maternal presence fosters the activity of this network in offsprings^4,12^. To evaluate the effect of maternal presence on the functional connectivity within the DMN network in foals we calculated the mean Pearson correlation coefficient between time-series of spontaneous variations of BOLD signal extracted from the homologous regions involved in DMN and DMN-like described in both humans and animals (aG, aCC/pFC and RSC). Using multiple regression analysis, we demonstrated that spontaneous BOLD signal fluctuations in these regions are indeed highly correlated (voxel-level threshold p < 0.001, t_(21)_ = 3.5272, FDR corrected, **Fig. 5c, left panel**), revealing the presence of a strong large brain network. Additionally, anticorrelated voxels were found bilaterally within the somatosensorial cortical area (S1), which is consistent with previous descriptions of DMN and DMN-like networks in humans, mice and rats. Consequently, we propose that altogether, the aG, aCC/pFC and RSC regions form a DMN-like network in domestic horse akin to what has been described in other mammals. To evaluate the DNM functioning in both experimental groups, we performed multiple regression analysis specifically focused on animals raised either with or without their mother (“*maternal presence*” group, **Fig. 5c, centre panel**). The analysis performed on animals raised with their mothers revealed cluster size and spatial organization of DMN like the one found in our population analysis. This result contrasts with the results obtained with animal raised without their mothers (“*maternal absence*” group, **Fig. 5c, right panel**) for which we found a smaller DMN and those a decreasing of the functional connectivity within the DMN in this population. These data suggest that presence of the mother could be involved in the maturation of the DMN-like brain network of offsprings in domestic horse.

### Prolonged maternal presence shapes behavioural development

Analyses of the percentage of scans spent in each behavioural activity combining all periods of observations (from the 1^st^ to the 7^th^ month of prolonged maternal presence) using *Aligned Ranks Transformation ANOVAs* showed that animals belonging to the “*maternal presence*” group displayed higher levels of exploration (p = 0.002), rest (p = 0.02), social interaction (for any type: p < 0.001; and for affiliative ones: p < 0.001) and lower time spent feeding (including suckling, p < 0.001) (**Fig. 6a**). For these animals, the percentage of scans showing suckling was 2.08±0.36% at 1-month post-weaning, 2.19±0.55% at 3 months post-weaning, and 2.83±0.70% at 7 months post-weaning (mean ± SE). No offsprings belonging to the “*maternal absence*” group were observed suckling. Social Network Analysis based on spatial proximity matrices (*i.e.* less than 1m between two animals) were used to calculate individual centrality scores^32^. Animals belonging to the “*maternal presence*” group had a higher centrality score (p = 0.002). Interestingly, foals did not differ in the percentage of time spent isolated (*i.e.* more than 10m from any other group member, p = 0.65). Behavioural tests were conducted at +7 months post-maternal separation to experimentally assess foals’ reactivity profiles in the social dimension (sociability test), non-social dimension (novel object test individually and novel surface test) and both dimensions together (novel object test in group). Results show that foals raised with their mother approach an unfamiliar horse more rapidly during the sociability test and required fewer training sessions to reach compliance for the care training (**Fig. 6b**).

**Fig. 6.**
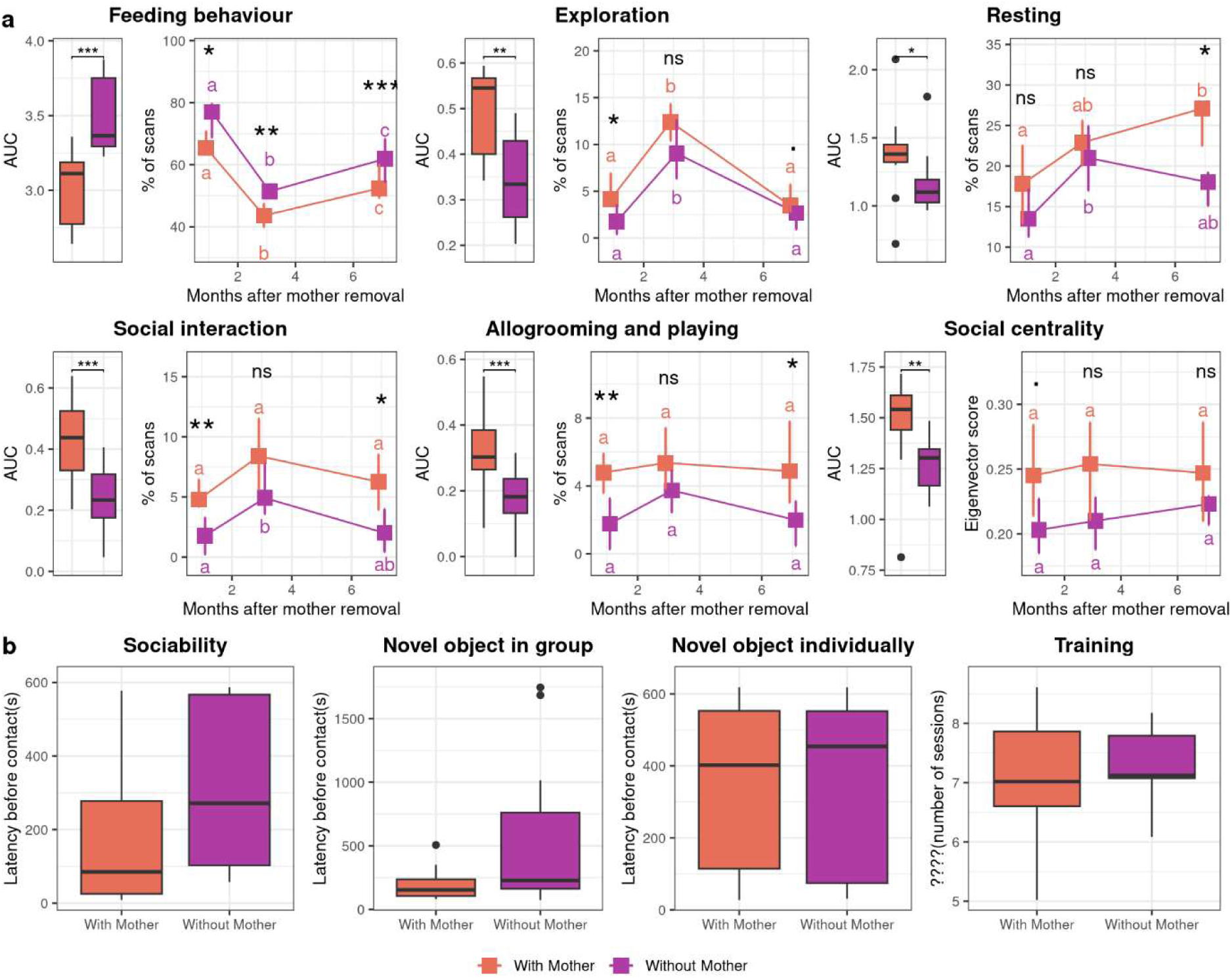
Prolonged maternal presence promotes sociability, exploration, and feeding efficiency. Longitudinal observations from +1 to +7 months post-weaning. Boxes represent the median, with error bars indicating the 95% confidence interval of the median (**a**). Medians sharing the same letter are not significantly different. Stars indicate significant differences between foals with or without their mothers (***: p<0.001, **: p<0.01, *: p<0.05, ns: non-significant). Behavioural tests conducted at +7 months post-weaning (**b**). The variables presented were selected based on multivariate analysis. Boxplot whiskers extend to ±1.5× the interquartile range for each treatment.

### Prolonged maternal presence shapes physiological development

Temporal evolution of the effect of prolonged presence of the mother on physiological parameters of the offspring were tested using *Aligned Ranks Transformation ANOVAs* and post-hoc tests (**Fig. 7**). Overall, animals belonging to the “*maternal presence*” group have a higher weight gain (p < 0.001), despite less time spent feeding, have higher concentrations of triglycerides (p = 0.02) and cholesterol (p = 0.002), and lower cortisol concentration (p = 0.002). The higher weight gain was observed at every time point (at +1 and +3 months: p < 0.001, at +7 months: p =0.002). The higher level of triglycerides in the “*maternal presence*” group was more pronounced at +1 month (p = 0.04) and +7 months since the beginning of treatment (p = 0.005), whereas the higher level of cholesterol was more pronounced at +1 month (p < 0.001), and +7 months (p = 0.01). Lower level of cortisol appeared to be restricted to the +3 months post-maternal separation time (p = 0.02). No differences have been observed for oxytocin, glycemia, IGF-1 and insulin.

**Fig 7.**
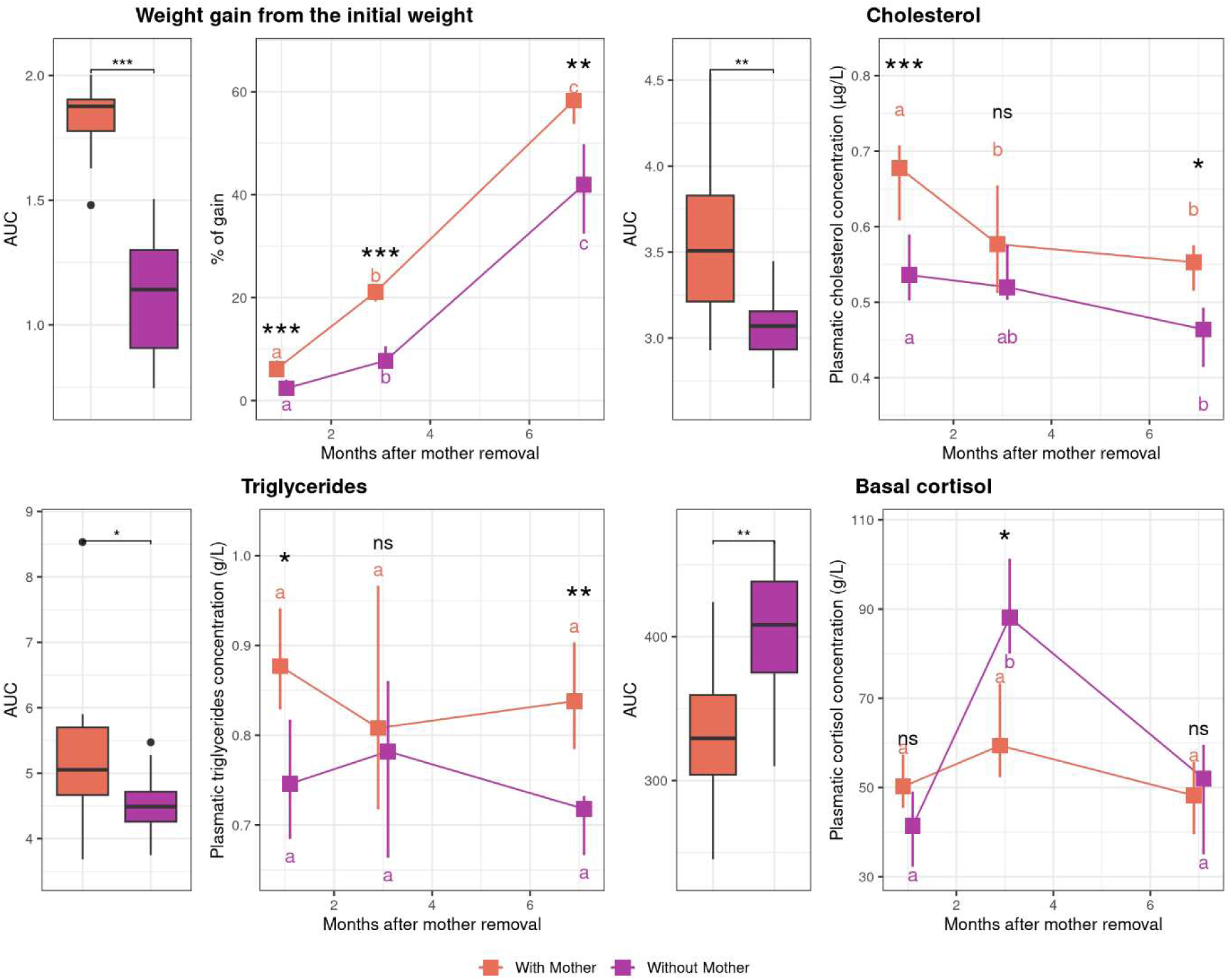
Prolonged maternal presence influences weight gain, basal cortisol, cholesterol and triglycerides plasmatic concentrations over time. Boxes indicate the median. Error bars indicate the 95% confidence interval of the median. Median sharing a letter are not significantly different. Stars represents significant differences between foals with or without their mothers (***: p<0.001, **: p<0.01, *: p<0.05, .: p<0.1, ns: non-significant).

### Relationships between brain structure and function and biobehavioural trajectories

The correlation structure among the six data groups (w*eight gain, non-social behaviour, non-social personality traits, sociality, MRI imaging*, and *physiology)* was investigated using a Multiblock Projection to Latent Structures models (PLS) with sparse Discriminant Analysis (multiblock PLS-sDA). This comprehensive multivariate approach optimises correlations across various data groups and conducts discriminant analysis to uncover a specific signature for each experimental group. The selected final model is presented in **Fig 8**. The hues and ellipses associated with the treatment demonstrate the discriminatory ability of each component to distinguish between presence and absence of the mother (**Fig. 8a**). While the centroids of each treatment are clearly separated, there exists a moderate degree of overlap between the sample groups within their confidence ellipses. Furthermore, the first components from each data group are correlated to at least one other group (indicated by the large correlation coefficients in the bottom left, **Fig. 8a**). Our final statistical model shows that the first components of variables for ‘*MRI imaging*’ are significantly correlated with ‘*sociality*’ (r=0.79), ‘*non-social behaviours*’ (r= 0.87), ‘*weight gain*’ (r= 0.87) and ‘*physiology*’ (r=0.9). Strikingly, a redundant pattern of antagonist correlations between the functional activity of aCC and RSC-Cu/PrCu (ALFF) has been found for +1-month social interaction and social isolation, +1/+7 months playing and allogrooming (*i.e.* affiliative interactions), +1/+3/+7 months feeding behaviours, +1/+3 months weight gain, +1/+7 months cholesterol and +7 months triglycerides. Additionally, the functional activity of *amygdala* (left and right – ALFF) is largely correlated with the same variable and in a similar direction than the functional activity of anterior cingulate cortex (aCC). Finally, microstructural modifications (VBM) within the mid-cingulate cortex (mCC), hippocampus and insula (left and right) are all negatively correlated with +1/+3 months weight gain and +7 months triglyceridemia (**Fig. 8b**). In conclusion, our model shows that animals belonging to the “*maternal presence*” group display higher functional activity within both the aCC and amygdala (left/right), are more social and less isolated up to 7 months post-treatment, whereas animals belonging to the “*maternal absence*” group display a higher functional activity of the RSC-Cu/PrCu, higher GMC values within hippocampus and mCC and spend more time feeding without gaining more weight (**Fig. 8c**).

**Fig 8.**
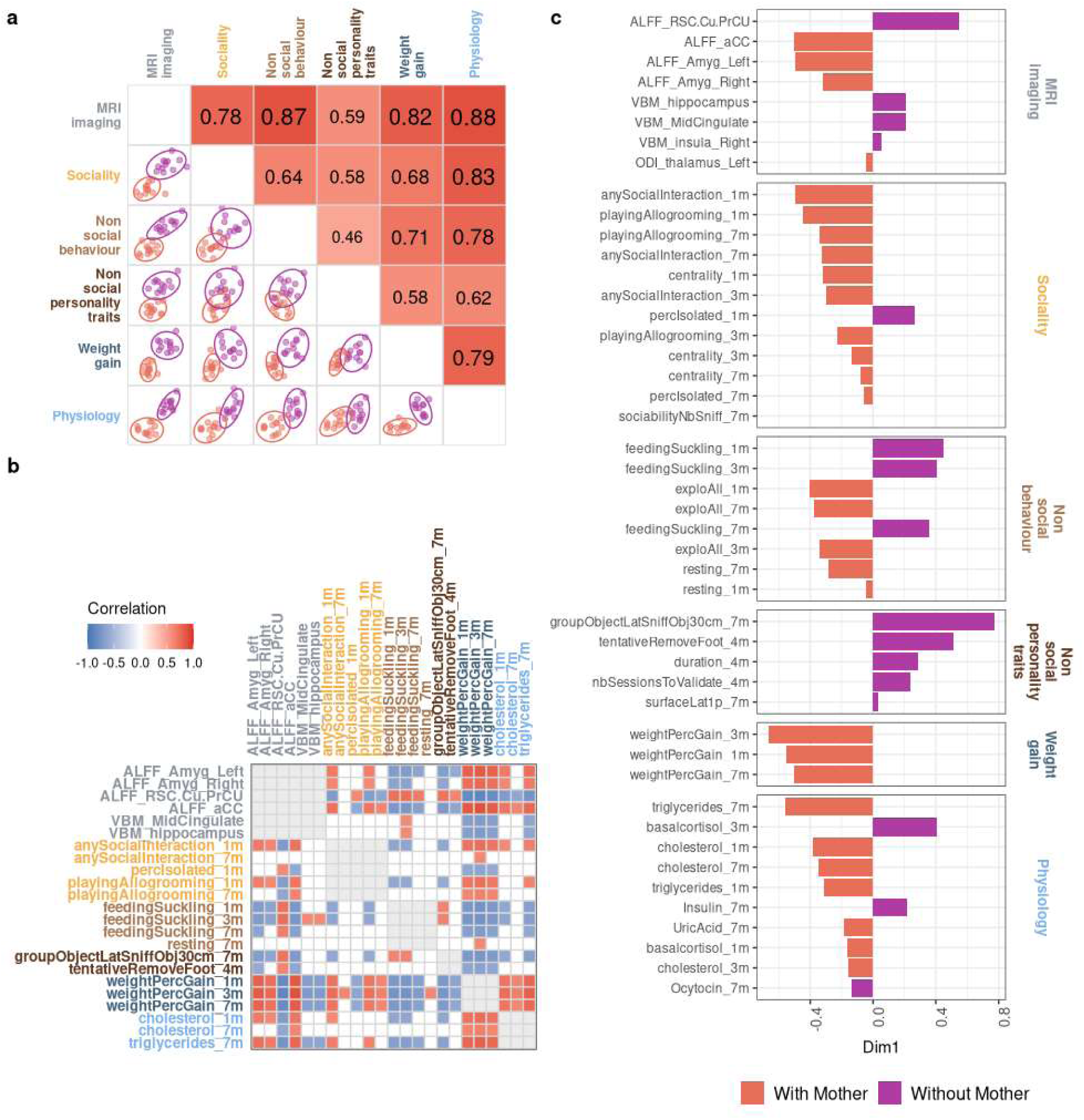
Multiblock sPLS-DA identifies differences between foals and correlations between variables. Correlation between the first dimensions of each group of variables (**a**). Correlation between variables belonging to two different groups (**b**). Loading weights of each selected variable on each dimension and each data group (**c**). The colour indicates the class in which the variable has the maximum level of expression using the median. The most important variables (according to the absolute value of their coefficients) are ordered from right to left.

## Discussion

Mapping the association between brain, behaviour and physiology is a critical objective in neuroscience. In this context, multivariate methods have gained popularity and Latent Structures models (PLS) emerged as tools to uncover complex brain-behaviour-physiology associations, providing a better understanding of how structural and functional brain organizations are linked with physiology and/or behaviour^33^. These advanced statistical approaches allowed us to combine brain imaging metrics with longitudinal biobehavioural assessment in foals and revealed that social affiliative behaviours during the mid-childhood fosters the development of the allostatic regulatory system in the offspring in domestic horse through two fundamental biological dimensions: 1/social life and feeding and 2/metabolic efficiency.

First, our results support those affiliative behaviours play a critical role in shaping social trajectories. During mid-childhood, offsprings’ functional brain metrics of RSC-Cu/PrCu and aCC/PFC, two major hubs of the default mode network (DMN), were shown by our statistical models to be influenced by social interactions with caregivers. Specifically, offsprings benefiting from prolonged maternal presence during this critical period exhibited lower RSC-Cu/PrCu activity but higher aCC/PFC activity. By comparison, human brain imaging studies reported distinct roles for these two DMN hubs in social behaviour. The RSC-Cu/PrCu region (PCC in humans) is primarily involved in self-other distinctions, while the aCC/PFC region is associated with monitoring both one’s own and others’ mental states. Moreover, the DMN has been found to play a critical role in self-reference, episodic and autobiographical memory, theory of mind, and other high-level social cognitive functions in humans^34^. While the DMN has been previously identified in a few animal species (e.g., mouse, rats, marmosets, chimpanzees)^35^, our study provides the first evidence of a DMN-like network in horses. Furthermore, our results highlight that this network, central for complex social behaviour in humans, may also support optimal social development in domestic horses. Indeed, our results also show a statistical association between the functional activity of these DMN regions and the foals’ social development. Offsprings benefiting from prolonged maternal presence exhibited more positive and adaptive social behaviours, alongside with increased exploratory and resting behaviours. Strikingly, these developmental advantages emerged despite all young individuals being raised in the same socially rich environment, highlighting the specific role of the mother-offspring bond in shaping these trajectories. This also underlines that the presence of adult models or peers alone is insufficient for optimal social development, as similarly observed in primates^1,2^. Taken together, these findings underscore the essential role of prolonged maternal presence in fostering successful development of social competences, and suggest that the DMN may serve as a founding structure for social development across social mammalian species^36^. The mother-offspring dyad thus provides a crucial social environment that fosters the maturation of DMN involved in the development of social abilities.

Second, our study suggests a caregiver-driven maturation supporting the physiological control of feeding and metabolic trajectories. Indeed, consistently throughout our study, foals benefiting from prolonged maternal presence spent less time feeding and displayed higher levels of exploration and rest compared to those separated from their mothers. Interestingly, they also showed higher weight gain and higher concentrations of triglycerides and cholesterol, consistent with previous findings in this species^7^. Because these physiological parameters are intimately linked to feeding efficiency, metabolism, and energy storage, these results highlight a certain advantage in the metabolic and growth status of foals benefiting from prolonged maternal presence. First, lipid metabolism is essential for the development of the central nervous system, as cholesterol is a primary component of neuronal membranes and intracellular structures. Numerous longitudinal and cross-sectional studies have linked low blood cholesterol and triglyceride levels to impaired cognition, recognition memory, and inhibitory processing in humans^37^. In the present work, offsprings with prolonged maternal presence preserved higher cholesterol and triglyceride levels and achieved better weight gain while spending less time feeding, suggesting that they maintained more efficient feeding behaviours and/or metabolic functioning. Several mechanisms could account for this metabolic efficiency. Access to maternal milk, although it represented only ∼2.5% of the foals’ total activity budget, may still have provided valuable nutrients beneficial for growth. The benefits of lactation are well established in humans as prolonged breastfeeding confers advantages for both physiological and cognitive development^38^. Despite both experimental groups consuming primarily solid diets, even minimal maternal lactation might have conferred some physiological benefits for these animals. Additionally, feeding is a goal-directed behaviour essential for the survival of all metazoan organisms which is believed to be rooted within innate neuronal circuits, particularly within the hypothalamus^39^. In our study, we found differences in the microstructural organization of the hypothalamus between the two groups, confirming that fundamental feeding pathways might be influenced by prolonged maternal presence. Moreover, social learning and emotional regulation could further refine feeding behaviour through additional, higher-order mechanisms. Indeed, we found that functional brain metrics in aCC/PFC and RSC-Cu/PrCu (DMN) were associated with feeding behaviour, suggesting a role for higher-order cognitive processes in regulating food intake. In line with this, research in humans has revealed DMN alterations in patients suffering from eating disorder (e.g. *anorexia nervosa*), linking the disruptions in cognitive control networks to feeding disorder^40^. Our statistical model also identified a significant association between amygdala functional activity and feeding behaviour, consistent with previous studies in mouse showing that specific neuronal populations in the central nucleus of the amygdala integrate negative affective states signals such as fear/anxiety and impacting feeding behaviours^41^. Prolonged maternal presence could limit these negative affective states through social buffering mechanisms, as suggested by lower cortisol levels observed after 3 months of prolonged maternal presence, ultimately supporting the development of more efficient feeding behaviours in offsprings. Taken together, these findings underline how a symphony of social, behavioural, neural, and physiological processes in foals with prolonged maternal presence leads to more efficient and optimal feeding and growth. This complex synergy likely reflects an allostatic regulatory process nurtured by a sustained maternal relationship, ultimately fostering healthier growth and neural development.

Ultimately, the present study allowed for the implementation of dedicated resources for *in vivo* MRI investigations of the horse brain. In the era of digital neuroscience and open science, preclinical MRI is a booming hot topic. However, the availability of MRI-compatible populational brain templates and atlases is mandatory for studying the brain of new animal models in neurosciences that the scientific community increasingly advocate for^42–44^. Together, templates and atlases allow group analysis, which addresses size and volume of pivotal brain structures and reveal both functional and structural brain networks^45^ as well as brain microstructural organization *in vivo*^28^. The multimodal abilities of MRI are powerful and its unique translatability between species enlarge the horizon of the field. Among animal models used in neuroscience, the domestic horse is of special interest due to its outstanding cognitive performances^46,47^. Indeed, horse exhibit testable cognitive function and are able to the perform learning, discrimination, match-to-sample and memory tasks^48^. Additionally, horse is the longest-lived of the common domestic species (25–30 years) making this animal model particularly relevant for the study of slow progressive neurological diseases^46,49^. Hence, in the first step of our study, we generated a new set of neuroinformatic tools offering an unprecedented and complete resource dedicated to MRI studies of the horse brain *in vivo* namely, a high-resolution brain template and its associated GM, WM and CSF priors built from 23 individuals for anatomical, functional and diffusion imaging analysis in horses. We also delivered the first *in vivo* horse brain atlas composed of a mosaic of 115 regions of interest which offers an accurate segmentation of both cortical and subcortical areas. This comprehensive set of MRI compatible template will allow standardization of horse brain MRI data analysis and paves the way for the development of multicentric preclinical studies in horses as previously delivered for rats^50^.

## Conclusion

Our study demonstrates that, even beyond the exclusive nursing phase, the mother-offspring relationship exerts a powerful influence on the development of offsprings’ brain, behaviour and physiology. Despite a socially enriched environment with peers and adult conspecifics, only offsprings benefiting from prolonged maternal presence displayed consistent advantages in social behaviour, feeding and metabolic development. This finding highlights the uniqueness of the maternal bond in fostering offspring brain maturation at an advanced developmental stage. We also reveal changes in key neural cortical networks and pivotal subcortical region such as the hypothalamus and amygdala and present here the first *in vivo* demonstration of a DMN-like functional network in horse. By coupling these discoveries with dedicated MRI-based resources, our study positions the horse as an original large-mammal model for investigating extended caregiver-offspring relationships. These insights also invite to reconsider early weaning practices for animals under human care, as preserving maternal contact may yield significant long-term benefits for both animal welfare and developmental trajectories.

## Methods

### Subjects

Subjects were 24 Welsh foals, 12 females and 12 males, aged 6.64 ± 0.07 months (mean ± SE) and their 24 mothers aged 8.2 ± 0.3 years on the day of the removal of the mothers. All animals (foals and mothers) were born and kept at the <u>Animal Physiology Experimental Unit</u> (UEPAO, INRAE, 37380 Nouzilly, France). They all lived in groups in, depending on the season, indoor collective stalls (20 m x 25 m) on straw with free access to an outdoor paddock (from November 26^th^, 2021 to April 23^th^, 2022), or in large outdoor grass pastures (April 23^th^, 2022 to June 27^th^, 2022). The dietary ration, primarily based on *ad libitum* access to forages, pasture and mineral block, was designed to meet the theoretical nutritional requirements (INRA 2011 tables) for weaned foals of this type and age^51^. Foals born between May 13^th^ and June 27^th^, 2021. During the first five months of life, they were all kept with their mothers in the same outdoor pasture. In October 2020, two distinct herds were formed and kept in two different pastures: 12 dyads of male foals/mothers (Herd A), and 12 dyads of female foals/mothers (Herd B). The protocol then started from November 2020 (**Fig 1a**). All procedures were performed in accordance with the European directive 2010/63/EU for animal protection and welfare used for scientific purposes and approved by the local ethical committee for animal experimentation (CEEA VdL, Tours, France, ref. APAFIS #32985-2021091516064302).

### “With mother” and “without mother” treatments

Half of the subjects were assigned to the “*maternal presence*” treatment (N=12: 6 males and 6 females) and half of them to the “*maternal absence*” treatment (N=12: 6 males and 6 females). The “*maternal presence*” and “*maternal absence*” foals were balanced according to date of birth, weight and paternal origins. The “*maternal absence*” treatment consisted in removing half of the mothers in each group (A and B) at the same time when the foals aged 6.64 ± 0.07 months old (mean ± SE). Consequently, after maternal separation, each herd (A and B) was composed of 6 without-mothers foals, 6 with-mothers foals and their 6 adult mothers. Therefore, presence or absence of the mother was the sole difference between with or without mothers’ foals. Foals have been kept within the same social herd (A or B) and could therefore interact with each other and with the adult mothers still present. One female foal belonging to the “with mother” leave the experiment 4-months after the maternal separation procedure due to medical reasons. Imaging, behavioural and physiological data from this animal have been retrieved for the final analysis.

### Magnetic Resonance Imaging

#### Anaesthesia

Imaging procedures have been realised in the <u>PIXANIM imaging facilities</u>, on the same site as animal’ housing. The complete procedure has been described within *Supplementary Material* and inspired from our previous study^52^. Briefly, anaesthesia was induced in two successive stages. First, a dose of romifidine hydrochloride was injected intravenously as a premedication. Five minutes afterward, ketamine hydrochloride was injected intravenously to induce anaesthesia. Animal was placed on supine position onto the MRI table and a saphenous catheter, and a urinary drain were then placed, and an ophthalmic ointment was applied to avoid ocular dryness. A veterinarian and an animal technician were constantly present to monitor animal’s respiratory and cardiac rate. The mean duration of anaesthesia from the induction to the last injection was approximately of 3h.

#### In vivo MRI acquisitions

MRI data have been acquired on 3 Tesla VERIO Siemens systems (Erlangen, Germany), using two flexible coil (Siemens FLEX Large 4 elements) tied around the head and three MR acquisitions were performed on each animal. MR sequences have been optimized to 1/ be performed in a time compatible with anaesthesia (approximately 3h), 2/ reduce artefacts (folding, truncation, etc.) and 3/ optimise SNR. Regarding these specifications, the following sequences were used:

- for brain morphometry investigations, a three dimensional T_1w_ MPRAGE acquired in coronal plane was used with the following parameters: Echo Time/Repetition Time=2.67ms/2500ms, Flip Angle=12°, Inversion Time=900ms, Number of Excitation=3, Partial Fourier=1, Slice Thickness=1 mm, Slice Number=208, Field of View=256×256mm, matrix=256×256, final resolution 1mm^3^.
- To investigate the brain microstructure, we used a diffusion weighted MRI protocol based on a two dimensional T_2w_ spin-echo sequence acquired in the axial plane, over 3 different shells optimized for Neurite Orientation Dispersion and Density Imaging (NODDI) model using the following parameters: Shell 1: b=300 s/mm^2^, 6 directions; Shell 2: b=700 s/mm^2^, 30 directions, and Shell 3: b=2000 s/mm^2^, 64 directions. The fixed parameters are Echo Time/Repetition Time=109ms/11.5s, Flip Angle=90°, Number of Excitation=1, Partial Fourier=0.75, Slice Thickness=2.4mm, Slice Number=57, Field of View =256×256mm, matrix=128×128, final resolution 2×2×2.4mm^3^, one b=0 per shell). Sequences have been acquired in different reading phases (left-right and right-left) for distortion corrections.
- To investigate brain functioning a T_2w_ spin-echo-planar imaging (SE-EPI) sequence acquired in the axial plane and left/right reading phase was used with the following parameters: Echo Time/Repetition Time=24ms/3.97s, Flip Angle=90°, Number of Excitation=1, Partial Fourier=1, Slice Thickness=3.3mm, Slice Number=40, Field of View=220×220mm, matrix=110×110, final resolution 2×2×3.3mm^3^, Number of repetition=250. Additionally, two similar sequences in different reading phases (left-right and right-left) of ten volumes each were acquired for distortion corrections. Only one case of myositis has been detected after the imaging procedure, nevertheless the animal fully recovered within a week. DICOM data were converted to NIFTI format and organized as standardized data sets according to the Brain Imaging Data Structure (BIDS) using <u>BIDScoin</u> and are downloadable on <u>Zenodo</u>.

#### TEBTA template creation

T_1w_ MPRAGE data acquired for each animal were noise and signal bias corrected using <u>Ginkgo</u> and <u>N4BiasFieldCorrection</u> respectively and coregistered to the Johnson *et al* template^46^ using <u>antsRegistrationSyNQuick</u>. Coregistrated data were used to segment each brain using <u>SPM</u> and the GM, WM and CSF priors provided by Johnson *et al.* to create the GM, WM and CSF probabilistic maps of each subject. Denoised and signal bias corrected images were used in parallel to create a study specific T_1w_ template, using <u>modelbuild</u>, an optimized pipeline using <u>antsMultivariateTemplateConstruction2</u>, an unbiased template building method developed in ANTs package. Once both linear and non-linear transformations were calculated for each animal, we compiled all the transformations calculated for each image and applied them once to corrected images using <u>antsApplyTransforms</u> to limit interpolation effects. Resulting images have been used to create the <u>TEBTA template</u> by calculating the mean image of each normalized T_1w_ using <u>Ginkgo</u>. Probabilistic maps (GM, WM, CSF) have been normalized using the previous linear and non-linear transformations and GM, WM and CSF priors have been created by calculating the mean image of each normalized map using <u>Ginkgo</u>. Both templates and priors will be used for spatial normalization and segmentation for VBM analysis (see *Voxel Based Morphometry Analysis* section). The complete procedure is described *Supplementary Material*.

#### TEBTA atlas creation

For the creation of the <u>TEBTA atlas</u>, the equine brain atlas provided by Johnson *et al* is used as a starting point. Firstly the Johnson’ template is linearly and nonlinearly coregistrated within the <u>TEBTA template</u> using <u>antsRegistrationSyNQuick</u>. Then, both linear and non-linear transformations were applied to each ROI of the Johnson’ atlas using <u>antsApplyTransforms</u>. Each normalized ROI has been visually inspected to check boundaries and accuracy of registration. Then some regions of interest (ROIs) such as *Corpus Callosum*, *arbor vitae*, and ventricular systems have been updated/added using WM and CSF priors for a best fit to these ROIs to the <u>TEBTA template</u>. Eventually, additional subcortical ROIs such as septum, preoptic hypothalamic area, nucleus accumbens, striatum, etc. have been drawn and delimited manually using <u>fsleyes</u> and <u>itksnap</u>. To propose a valuable parcellation of the equine cortex, we implemented the methods previously used by Garin *et al* for the establishment of a functional atlas in lemur mouse^53^. Briefly, a multi-animal dictionary learning statistical analysis was performed with Nilearn (random_state = 0) on preprocessed rsfMR images (see *Functional Imaging Analysis* section)^54^. A mask excluding the WM, CSF and subcortical areas was used to restrict the dictionary learning analysis to cortical functional data. The study based on 60 sparse components was selected for the final analysis. Each bilateral component was split into two unilateral regions and labelled left or right. This led to a 3D functional atlas composed by a mosaic of 55 local functional regions that were named using <u>itksnap</u>. The name of each ROI was defined using the names of brain structures reported by Schmidt *et al*^55^, the AAL2 human brain atlas but also using their structural connectivity (see *TEBTA fiber atlas creation* section). The complete procedure is described *supplementary material*.

#### TEBTA fiber atlas creation: Structural connectivity of the equine brain

Cortical ROIs resulting from the dictionary learning segmentation of the equine cortex were used to identify the largest bundle tracts of white matter to help for the identification of ROI based on their structural connectivity. For the construction of the fibre tractogram, we used the analytical Q-ball reconstruction model and streamline regularized deterministic (SRD) tractography algorithm available in <u>Ginkgo</u>^56^ (see details in *supplementary methods*). From obtained populational tractogram, we selected the large bundles (length>150mm) using a 2 steps approach: 1/ bundles between each cortical ROIs have been selected by a ROI-to-ROI selection, labelled and merged. Then bundles between thalamus and each cortical ROI have been selected as well as bundles between cerebellum and each cortical ROIs. With this selection, we expected to find long cortico-cortical pathways (i.e. *cingulum*), thalamic projections such as the somatosensorial tract to identify the somatosensory areas and the cerebellocortical tract to identify the motor areas. To eliminate non-relevant fibres previously selected within the bundle tract was filtered by length (>150mm) and by tortuosity (<2σ of mean of tortuosity). 2/ bundles selected have been visually inspected and 7 large bundle tracts (cingulum, corticospinal tract, anterior/posterior/inferior thalamic radiations, inferior longitudinal tracts and cerebellocortical tract) have been identified on basis of morphology, position within the brain, location and by comparison of fibre atlases available in humans and NHP. Areas connected to these tracts have been named according to their structural connectivity, position and literature^55^. The complete procedure is described *supplementary material*.

#### Voxel-Based Morphometry Analysis

For voxel-based morphometry analysis (VBM), we used T_1w_ images. Data were first denoised and signal bias corrected using <u>Ginkgo</u> and <u>N4BiasFieldCorrection</u> respectively and linearly coregistered to the TEBTA template using <u>antsRegistrationSyNQuick</u>. Coregistrated data were then pre-processed with <u>SPM</u>. Each image was segmented into probability maps of GM, WM, and CSF using the default settings in the <u>SPM8 toolbox</u> and the <u>TEBTA</u> version of GM, WM, and CSF probability maps. Transformation matrices obtained were used to normalize GM, WM and CSF probability maps obtained for each subject. GM and WM probability maps of each scanning session were normalized to our stereotaxic space using transformations matrices obtained herein and resampled. Normalized GM and WM images were used for diffeomorphic anatomical registration using exponentiated lie algebra (DARTEL) to calculate diffeomorphic flow fields. Each normalized GM image was then warped using deformation parameters calculated by the DARTEL routine and then modulated to correct the volume changes that may have occurred during the deformation step. Finally, normalized-warped-modulated GM images were spatially smoothed by convolving with a 4 mm full width at half-maximum (FWHM) isotropic Gaussian kernel to create GMC maps. To assess regional GM changes over the groups (“*maternal presence*” n = 11 *versus* “*maternal absence*” n = 12), GMC maps were compared using an unpaired Student t test proposed by <u>SPM</u>. A brain mask was used to constrain the analysis to the brain. For each cluster, the significance of the peak voxel was set as p < 0.05 (t-score = 1.72, degree of freedom=21). The results are presented on an axial and sagittal brain slice series generated with <u>Nilearn</u>.

#### Diffusion imaging data analysis

Because of its practical implementation, the NODDI model has become very popular to map the tissue microstructure *in vivo* and *ex vivo* in both clinical^57,58^ and preclinical applications^59^. The NODDI model relies on a biophysical model that separates the diffusion of water into three diffusive compartments (intra-neurite, extraneurite and CSF)^28,57^, which are supposed as non-exchanging, contributing to the global diffusion attenuation. The net diffusion signal attenuation (A) corresponds to the following linear combination *A* resulting from a linear combination of the individual signal attenuations associated with each compartment, including: 1/ the neurite compartment of water molecules trapped within axons and dendrites characterized by a volume fraction (fic), 2/ the extra-neurite compartment characterized by a volume fraction (fec) and 3/ the CSF compartment containing free molecules with an isotropic displacement probability characterized by a volume fraction (fiso). Hence, the net signal diffusion signal A corresponds to the following linear combination:

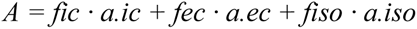

We pre-processed raw multishell diffusion imaging data and mapped estimated each of the previously described fractions using <u>Ginkgo</u>. First multishell diffusion imaging data were corrected for magnetic susceptibility distribution (<u>topup</u>), motions and eddy current-induced distortions (<u>eddy</u>). Then mean of B0 data for each animal was calculated and resulting images were used to calculate a diffusion weighted template using by <u>modelbuild</u>. Resulting template was then normalized to the <u>TEBTA template</u> using <u>antsRegistrationSyNQuick</u>. Then <u>Ginkgo</u> was used for NODDI analysis (<u>DwiMicrostructureField</u>) and released 4 maps: the fractional anisotropy, the intraneuritic fraction, the orientation dispersion index, and the isotropic fraction. Both linear and nonlinear transformation calculated by <u>modelbuild</u> and <u>antsRegistrationSyNQuick</u> were applied once to each contrast for spatial normalisation of the data within the TEBTA space using <u>antsApplyTransforms</u>. Then, images were spatially smoothed with an isotropic Gaussian kernel by convolving a 4 mm full width at half maximum. To assess regional FA, ICF, ODI and isoF changes over the groups each contrast was compared using an unpaired Student t test proposed by <u>SPM</u>. A brain mask was used to constrain the analysis to brain. For each cluster, the significance of the peak voxel was set as p < 0.05 (t-score = 1.72, degree of freedom=21). The results are presented on an axial and sagittal brain slice series generated with <u>Nilearn</u>.

#### Functional imaging data analysis

rs-fMRI data were pre-processed as previously described^24,25,45^. Briefly, EPI images were corrected for slice timing (<u>slicetimer</u>), motions (<u>antsMotionCorr</u>) and susceptibility distribution (<u>topup</u>). Then mean of EPI data for each animal was calculated and resulting images were used to calculate a functional template using by <u>modelbuild</u>. Resulting template was then normalized to the <u>TEBTA template</u> using <u>antsRegistrationSyNQuick</u>. Then images were detrended, low- and high-pass filtered (0.01Hz – 0.01Hz) the effect of the six previously calculated motion parameters including translations and rotations, both WM and CSF and global signals, were removed from data through linear regression using <u>Nilearn.</u>

#### Fractional amplitude of low-frequency fluctuations analysis

The fractional amplitude of low-frequency fluctuation (0.01–0.08 Hz) values were computed using the previously processed 4D data the <u>fALFF function</u> from <u>1000 Functional Connectomes Project</u> was used to reveal the temporal and regional changes in GM occurring in fALFF maps. Then fALFF maps were spatially normalized to the <u>TEBTA template</u> using both linear and nonlinear transformation calculated previously and applied once using <u>antsApplyTransforms</u>. To assess regional fALFF changes over the groups each contrast was compared using an unpaired Student t test proposed by <u>SPM</u>. A brain mask was used to constrain the analysis to the brain. For each cluster, the significance of the peak voxel was set as p < 0.05 (t-score = 1.72, degree of freedom=21). The results are presented on an axial and sagittal brain slice series generated with <u>Nilearn</u>.

#### Identification of the equine Default Mode Network (DMN)

To elicit for the first time the DMN in equidae, we calculated the mean Pearson correlation coefficient between time-series of spontaneous variations of BOLD signal, extracted from the functional ROIs previously calculated by the dictionary learning approach we defined functionally homologous to regions involved in DMN and DMN-like described in humans in numerous species (aG, aCC/pFC and RSC). Using data extracted and a multiple regression analysis approach we demonstrate that spontaneous BOLD signal fluctuation in these regions is highly correlated. For each cluster, the significance of the peak voxel was set as p < 0.001 (t-score = 3.5272, degree of freedom=21, FDR corrected). The results are presented on an axial and sagittal brain slice series generated with <u>Nilearn</u>.

### Behavioural observations

Behavioural observations occurred during the day between 10:00am and 3:30pm. For each observational period (+1 month, +3 months, and +7 months post-maternal separation), foals were observed across the same week, for two to five days. For each period, foals were therefore observed during 10 to 20 hours in total. Behavioural observations consisted in scan samplings^60^. Every 15 min, for each foal, its activity, if it is isolated or near another animal (≤10m), the identity of its closest neighbour when relevant and the distance between them in metres. A total of 40 to 84 scans/period/foal was therefore analysed. The proportion of scans spent in each of these activity categories has been calculated to estimate the time the foals spent: feeding (drinking and suckling included), exploring the environment, resting (standing or lying down), and interacting positively and actively (allogrooming or playing socially). In addition, an “isolation” score has been calculated based on the proportion of scans spent isolated (+10m away from another groupmate). The higher the time spent isolated, the less gregarious and the more socially independent the foal is. Finally, a centrality score, or eigenvector, has been calculated based on spatial proximities with the closest neighbour^61^. Two individuals were considered close when the inter-individual distance was less than 1m (i.e. distance enabling the individuals to interact without having to move). The eigenvector was obtained using SOCPROG 2.9^61^ using a two-entries proximity matrix. The eigenvector represents the total number of connections an individual has, including the connections between its partners. The higher this index is, the more central the individual is within its social group. All these parameters have been calculated for each foal for each period of observation (+1 month, +3 months, and +7 months post-maternal separation).

### Behavioural tests

All tests, apart from the social isolation test, were conducted in the foals’ home environment. Each test was carried out on a separate day within the same week of the 7^th^ month post-maternal separation period. In the morning of each test day, the entire group (i.e. mothers, “*maternal presence*” foals, and “*maternal absence*” foals, were assembled and confined in half of the collective stable, using removable tubular barriers and gates. This arrangement ensured that the test area was adjacent to the group area and familiar to the foals, thereby minimising any potential novelty effect or social isolation. Each individual test started in a standardised manner. A full opaque wooden panel (dimensions: height 2m, length 4m) was installed to create a specific corner in the test area that remained hidden from the rest of the group. This ensured that the stimulus (such as an unfamiliar horse or object) was only visible and accessible to foals undergoing testing at that moment. Two experimenters entered the group area and approached the animal selected for testing, with the order of selection being semi-random and varied for each test session. The foal was then fitted with a head collar, with one experimenter holding a lunge line and the other experimenter walking behind the foal and guiding it with their arms opened. Together, they led the foal calmly to the test area through the gate separating the two sections. Upon reaching the test area, foal was released approximately 5m past the gate, positioned at a reasonable distance (20m) from the designated ‘stimulus’ area. This setup allowed the test to start, giving the foal the freedom to either approach the stimulus or remain away (see **Fig. S2** in *Supplementary Material*).

#### Sociability test

Before the beginning of the test, an unfamiliar horse (a female Welsh pony, 23 years old) was brought to the test area. Selected for its calmness and lack of reactivity to social isolation, the unfamiliar horse was tied up using its head collar and provided with forage and a bucket of water to ensure it remained relatively motionless, positioned parallel to the test area. The panel separating the unfamiliar horse, and the test foals consisted in a feed-through panel (*i.e.* an ensemble of vertical metallic bars from the floor to the top allowing the foal to pass the head and neck through the panel but nothing more). With this arrangement, the foal was allowed to approach and enter in physical contact with the unfamiliar horse using its head only. This setup permitted to foal to approach and make physical contact with the unfamiliar horse solely using its head, while still allowing the unfamiliar horse to step away if desired. Throughout the procedure, the unfamiliar horse remained calm and accessible to all test foals. Interactions between animals were mainly neutral, characterised by reciprocal exploration, or no interaction occurred at all. Test duration was set for 10 min, starting with the release of the test foal from its initial position. Latency before the tested foal contacted the unfamiliar horse was recorded, with a maximum score of 601 seconds assigned if contact did not occur before the end of the test.

#### Fearfulness test: novel object test (individually)

Prior to testing, an unfamiliar object (a pink fitness ball, diameter: 65 cm, see *Supplementary Material*) was positioned in the test area, away from the view of the group. Tested foal was then brought to the test area following the same procedure previously described and was released into the area for a duration of 10 min. During this time, the latency before the foal contacted the unfamiliar object was recorded. A maximum score of 601 seconds was assigned if contact did not occur before the end of the test.

#### Fearfulness test: novel surface test (individually)

Following the procedure previously described^62^, an unfamiliar plastic tarp (dim:3m x 3m) with a bucket containing pellets placed on top (bucket positioned centrally) was positioned prior to testing, away from the sight of the group. For this test, the test foal was led 5m away from the surface by one experimenter. Once in place, a second experimenter held the bucket and brought it to the foal, allowing it to ingest a mouthful of pellets. Then, the second experimenter would walk back to the surface, ensuring the foal remained attentive, and then return the bucket at the centre. Then foal was released by the first experimenter. With this setup, the foal was required to approach the tarp and place its two forelegs on it to access and consume the pellets. The test concluded either when the foal completed this action or after 6 minutes if the foal did not step on the surface at all. The latency before the foal stepped on the surface for the first time was recorded. A maximum score of 361 seconds was assigned if the foal did not step on the surface before the test concluded.

#### Fearfulness test in social context: novel object test in group

The novel object was a foldable tunnel covered by plastic fabric (diam: 0.6m, height: 4m, see *Supplementary Material*), commonly used for dog agility practice. Two experimenters led all the 11 or 12 foals from a same social group to the test area simultaneously, while the mothers remained in the initial group area. Upon arrival of all the foals in the test area, the novel object was unfolded and immediately suspended 80cm above the floor using a sliding string connected to the ceiling of the building. This string was actioned by a third experimenter, ensuring the object was visible to all present animals from the same moment. Once the object was in place, all animals were allowed to move and explore freely for a period of 30min. During this time, the latency before each foal contacted the unfamiliar object was recorded. A maximum score of 1801 seconds was assigned if contact did not occur before the end of the test.

#### Social isolation test

This test, adapted from previous studies^63^, took place during the 4^th^ month following maternal separation. Each foal was individually led to a separate building located 150m away from their home stable. This was achieved by using a head collar and a rope, with one experimenter leading the foal and another providing guidance from behind. After arriving, each foal was promptly placed in an individual stable (4.5m x 3.5m) with the lower part of the door closed for 90s. From this position, the foal could see the experimenters but not any foals. It could hear sounds from foals that were not part of our study. The number of vocalisations (neighing) made by each foal during the test was recorded.

#### Standardized training and observation of care compliance in foals

Over a period of four consecutive weeks during the 5^th^ month post-treatment, we standardized and observed the routine hoof care training that all foals at the experimental unit undergo for health purpose. The training aimed to familiarize the foals with being touched on their limbs and voluntarily lifting their hooves, using a cooperation-based approach with food rewards. Hoof handling training was carried out to ensure the animals would tolerate routine farriery. We recorded the number of sessions required for each foal to meet predefined criteria for hoof lifting acceptance, as a proxy to assess the duration of training needed to achieve compliance. During the final week, we observed the farriery session, which was a necessary procedure for all animals. During this session, we also recorded the number of attempts each foal made to withdraw its hoof under standardized conditions.

### Physiological assessment

Blood samples were collected from all foals between 10:00 AM and 12:00 PM to avoid circadian variation, followed immediately by weighing each animal under constituent conditions and without restraint. Sampling occurred once prior to the maternal separation treatment and then at +1, +3, and +7-months post-separation. All samples were taken in the foals’ home collective stable, where the animals were handled calmly without the need for additional equipment, as this procedure was routine for them. Per sampling day, 20mL of blood was collected per foal, and then centrifuged to extract plasma that was then kept at −20°C until analysis. At the end of the experiment, plasma concentrations of cortisol, oxytocin, glycemia, cholesterol, IGF-1, and insulin were then measured to capture baseline levels of these physiological markers (see *Supplementary Material*).

### Data analysis

All statistical analyses were performed with R (version 4.2.2)^64^. Two foals have missing values: one for long-term behavioural tests and the other for MRI imaging data. To address this, missing data are imputed using Multiple Factor Analysis (MFA), using the imputeMFA function within the missMDA package^65^. Additionally, given that the foals were reared in same-sex groups, any potential effect of sex is mitigated using the ComBat method from the sva package^66^. This method enables the adjustment for batch effects in datasets where the batch covariate, in this case, sex, is known, employing an empirical Bayes framework. On these imputed and corrected data, a Multiple Factor Analysis (MFA) is performed on all the parameters using the FactoMineR R package^67,68^ to explore differences between foals with or without their mothers. Unlike Principal Component Analysis (PCA), MFA organises variables into groups, each carrying equal weight in the analysis. In this study, the six defined groups include *weight gain*, *non-social behaviour*, *non-social personality traits*, *sociality*, *MRI imaging* and *physiology*. Then, the correlation structure among the data groups was investigated using the mixOmics R package employing the DIABLO framework (Data Integration Analysis for Biomarker discovery using Latent cOmponents)^69,70^. This comprehensive multivariate approach optimises correlations across various data types and conducts discriminant analysis to uncover a specific signature to the foal groups. The underlying method used is a Multiblock Projection to Latent Structures models (MP-PLS) with sparse Discriminant Analysis (sDA). PLS components, which are linear combinations of variables, are constructed to maximise the sum of covariances across all blocks (i.e. groups of data). To choose the number of components, the function perf() is run with 3-fold cross-validation repeated fifty times. Examining the performance plot, we note an increase in both the overall and balanced error rates (BER) as the number of components increases from 1 to 2. All distance measures appear to yield similar accuracy, leading us to opt for the centroids distance. Based on this distance metric and the BER, we decide to set the number of components to 1. Furthermore, the weights of all pairwise covariances are determined by the design matrix. In this study and with the help of this analysis, we are seeking to determine the links between the variables of the different blocks. Consequently, we opted to establish the design matrix value at 0.8 for all block pairings, facilitating a thorough exploration of these associations. By employing the sparse option alongside discriminant analysis, the most discriminant variables between foal treatments are selected from each group of data. We choose the optimal number of variables to select in each data group using the tune.block.splsda() function, for a grid of keepX values for each type of data. This function has been set to favour a relatively small signature while allowing us to obtain enough variables for downstream interpretation. The function tune.block.splsda() is run with 3-fold cross validation and repeated fifty times. As this function is time-consuming, the grid of values tested for each data group has been reduced with a smaller number of repetitions. Subsequently, individual tests were conducted on kinetic variables identified through the preceding multivariate analysis. In order to examine the interaction effect between treatment and time, while also incorporating a random effect of individuals into the model, Aligned Ranks Transformation ANOVAs were performed^71^. Appropriate post-hoc comparisons were conducted to assess the differences in treatment at each time point, as well as the temporal evolution of each measurement^72^. These analyses were conducted using the R package ARTool. Same tests were applied to the area under the curve of each kinetic to obtain an overall difference. All adjusted p-values below 0.05 were considered as significant.

## Data availability

Datasets supporting the findings of this study along with the detailed statistical analyses are available are publicly available at https://zenodo.org/records/13969827. The Turone Equine Brain templates and atlas toolkit is publicly available at https://zenodo.org/records/10731031

## Code availability

All code used for magnetic resonance imaging (functional, voxel-based morphometry and diffusion) analysis are publicly available at https://github.com/DavidBarriere/Equisobrain. All code used for statistical analysis are publicly available at https://zenodo.org/records/13969827.

## Acknowledgements

M.V. and M.K. gratefully acknowledge support from the European HORIZON 2020 Marie Skłodowska-Curie Actions (MSCA) (project number: 101033271, MSCA European Individual Fellowship). All authors also thank the *Institut Français du Cheval et de l’Equitation* (IFCE) for funding this project. This work was also supported by the French National Research Institute for Agriculture, Food and the Environment (INRAE), which, through its specific experimental units <u>Unité</u> <u>Expérimentale de Physiologie Animale de l’Orfrasière</u>, the <u>PIXANIM imaging platform</u> belonging to the <u>UMR Physiologie de la Reproduction & des Comportements</u>, provided both animals and technical facilities for successful MR imaging on domestic horses. This work also benefited from the Phenotyping-Endocrinology laboratory for the hormonal assays and the IT infrastructure of the ISLANDe platform, and particularly a computing cluster financed by the European Regional Development Fund n°159037. The authors warmly thank the whole “equine” team of the Animal Experimental Unit for their technical support, and consistent commitment through the data collection and animal care, and especially Thierry Blard and Melinda Rousseau, along with Tiphaine Aguirre-Lavin. The authors are deeply grateful to the whole team of the Veterinary Clinic of the Nouvetière (Sonzay, France). Finally, the authors thank B. Piégu, for computer system administration and for help with ANTs-cluster deployment, and L. Croizier for her valuable help within behavioural data collection.

## Author contribution

Conceptualization: MV, FR, LC, LL, MK, DAB. Methodology development: MV, FR, HA, FE, IL, CD, CP, IU, DAB. Data collection: MV, FR, AG, HA, FE, JD, ALL, PB, YG, DAB. Project administration: MV, MK. Data validation and analysis: MV, GL, DAB. Writing of the manuscript: MV, GL, DAB. Commenting and editing the manuscript: HD, JD, LC, LL, ALL, HA, MK.

## Notes

https://zenodo.org/records/13969827

https://zenodo.org/records/10731031

https://github.com/DavidBarriere/Equisobrain

### Competing Interest Statement

The authors have declared no competing interest.

