## Supplementary Material for "Affiliative behaviours regulate allostasis development and shape biobehavioural trajectories in horse"

*Coauthors

Authors correspondence:

, and

**Supplementary Information**

### **Supplementary information – Brain Imaging**

The availability of *in vivo* population average templates and well-defined brain atlases is still a significant gamechanger for the use of the equine model in preclinical imaging^1–4^. Brain templates and atlas allow spatial normalisation and statistical analyses through group studies to compare the size/volume/fractions of pivotal structures based on microstructural imaging (anatomical MRI, diffusion MRI). Additionally, in the context of brain functional connectivity studies (fMRI) templates and cortical atlases allow the study of functional brain networks in different conditions^1–4^. To study equine brain using an MRI approach, only one resource was available^8^. This pioneer work proposes an *ex vivo,* unfixed, stereotaxic population template of the equine brain built from nine animals aged between 18 months and 30 years (7 males, 2 females) from different breeds (2 Quarter horse, 1 Mustang, 2 Standardbred, 3 Warmblood, 1 unspecified), three class of tissue segmentation maps for white matter (WM), gray matter (GM) and cerebrospinal fluid (CSF) and an atlas of 16 subcortical regions of interest^8^. However, this resource was characterised by significant limitations. First, all the animals used were adults at time of scanning and both gross and fine brain morphologies could be drastically different to our dataset in juvenile animals. Second, the cohort used to create both templates is sex-biased (~80% male *versus* 20% female) which is a limitation regarding the estimation of volumes in sexually dimorphic structures. Finally, the brain atlas proposed is largely incomplete with no cortical parcellation limiting, consequently the interest of such an atlas. In this context we built a new resource for the analysis of the equine brain, the [Turone Equine Brain Template and Atlas (TEBTA)](https://zenodo.org/records/10731031), a new set of standardised MRI compatible templates and atlas in stereotaxic coordinates meant to support the analysis of multimodal MRI data of the equine brain.

### **Supplementary methods – Brain Imaging**

#### **Imaging acquisition and construction of the TEBTA**

##### *Anaesthesia*

Imaging procedures have been realised in the [PIXANIM imaging facilities](https://eng-pixanim.val-de-loire.hub.inrae.fr/), on the same site as animal’ housing. Before being transported by van at the MRI platform (>5min course), animals were sedated to prevent stress (intramuscular injection of romifidine chlorhydrate 10mg/ml, single dose of 0.5mL Sedivet; Boehringer Ingelheim Vetmedica Inc, Beaucouze, France). Once at the MRI platform, anaesthesia was induced with a two steps process. First, an additional dose of romifidine hydrochloride 10mg/ml was injected intravenously as a premedication (0.14mg/kg, Sedivet). Five minutes afterward, ketamine hydrochloride 100mg/mL was injected intravenously to induce anaesthesia (2mg/kg, Imalgene, Boehringer Ingelheim Vetmedica Inc). Animal was then bedded in lateral recumbency and placed on a wheeled lift table. It was then placed on supine position onto the MRI table and maintained in this position with straps and memory foam mat and cushions to ensure stability, immobility and to prevent inadequate body pressure. A saphenous catheter and a urinary drain were then placed, and an ophthalmic ointment was applied on both eyes which were maintained closed during the whole imaging procedure to avoid ocular dryness. A veterinarian and an animal technician were constantly present near the animal to monitor the animal’s respiratory and cardiac activities and general state, and to ensure constant and conformable anaesthesia. Additionally, a camera zooming on the head of the animal in the MRI tunnel enabled additional monitoring from the control room.

During the functional imaging procedure (15min approximately), guaifenesin 100 mg/mL (Myorelax, Dechra Veterinary Products SAS, Montigny-Le-Bretonneux, France) by perfusion and a ketamine/ xylazine was administered by additional intravenously injections to ensure anaesthesia. At the end of the functional imaging procedure, small injections of ketamine/Xylazine and, additionally, diazepam (Valium®) were made regularly at the discretion of the veterinarian. The mean duration of anaesthesia from the induction to the last injection was approximately of 3h and animals received a mean volume of 528.57±31.54mL of the perfusion mix (containing guaifenesin), plus 18.74±0.003mL of ketamine, 2.64±0.17mL of diazepam and 2.70±0.10mL of romifidine chlorhydrate through injections (initial sedation and induction included).

##### *In vivo MRI acquisitions*

MRI data have been acquired on 3 Tesla VERIO Siemens systems (Erlangen, Germany), using a two flexible coil (Siemens FLEX Large 4 elements) tied around the head and three MR acquisitions were performed on each animal. MR sequences have been optimized to 1/ be performed in a time compatible with anaesthesia (approximately 3h), 2/ reduce artefacts (folding, truncation, etc.) and 3/ optimise SNR. Regarding these specifications, the following sequences were used:

- for brain morphometry investigations, a three dimensional T_1w_ MPRAGE acquired in the coronal plane was used with the following parameters: Echo Time/Repetition Time=2.67 ms/2500 ms, Flip Angle=12°, Inversion Time=900 ms, Number of Excitation=3, Partial Fourier=1, Slice Thickness=1 mm, Slice Number=208, Field of View=256x256 mm_,_ matrix=256x256, final resolution 1mm^3^.

- To investigate the brain microstructure, we developed a diffusion weighted MRI protocol based on a two dimensional T_2w_ spin-echo sequence acquired in the axial plane, over 3 different shells optimized for Neurite Orientation Dispersion and Density Imaging (NODDI) model using the following parameters: Shell 1: b=300 s/mm^2^, 6 directions; Shell 2: b=700 s/mm^2^, 30 directions, and Shell 3: b=2000 s/mm^2^, 64 directions. The fixed parameters are Echo Time/Repetition Time=109ms/11.5s, Flip Angle=90°, Number of Excitation=1, Partial Fourier=0.75, Slice Thickness=2.4 mm, Slice Number=57, Field of View =256x256 mm_,_ matrix=128x128, final resolution 2x2x2.4 mm^3^, one b=0 per shell). Sequences have been acquired in different reading phases (left-right and right-left) for distortion corrections.

-To investigate brain functioning a T_2w_ spin-echo-planar imaging (SE-EPI) sequence acquired in the axial plane and left/right reading phase was used with the following parameters: Echo Time/Repetition Time=24 ms/3.97 s, Flip Angle=90°, Number of Excitation=1, Partial Fourier=1, Slice Thickness=3.3 mm, Slice Number=40, Field of View=220x220 mm_,_ matrix=110x110, final resolution 2x2x3.3 mm^3^, Number of repetition=250. Additionally, two similar sequences in different reading phases (left-right and right-left) of ten volumes each were acquired for distortion corrections.

Once the imaging procedure finished, the animal was immediately transported back to a recovery stable to wake up progressively. Hydration (3 mL of Ringer minimum), flunixine meglumine (single dose of 3.5mL, Antalzen, 50 mg/mL, laboratorios calier, Barcelona, Spain) and of acefylline heptaminol (a single dose of 25mL of Vetecardiol, Intervet MSD Santé animale, Beaucouzé, France) were systematically administered. Veterinarian and at least two animal technicians were constantly present near the animal to monitor its recovery until it was able to walk steadily (approximately 1h40 for recovery). Then, the animal stayed in the recovery stable at least 6 hours after he stood-up for surveillance with another animal (the mother or another foal depending on the group) to avoid social isolation stress and finally joined back their housing. A blood sample was taken 48h after anaesthesia and sent to the veterinary office to check the absence/presence of myositis. Only one case of myositis has been detected afterward nevertheless the animal fully recovered within a week. DICOM data from the scanner were converted to NIFTI format and organized as standardized data sets according to the Brain Imaging Data Structure (BIDS) using [BIDScoin](https://bidscoin.readthedocs.io/en/stable/) and are downloadable on [Zenodo](https://zenodo.org/records/13969827).

##### *TEBTA template creation*

T_1w_ MPRAGE data acquired for each animal (23 scans) were noise and signal bias corrected using [Ginkgo](https://framagit.org/cpoupon/gkg) and [N4BiasFieldCorrection](https://github.com/ANTsX/ANTs/wiki/N4BiasFieldCorrection) respectively and coregistered to the Johnson *et al* template^8^ using [antsRegistrationSyNQuick](https://github.com/ANTsX/ANTs/blob/master/Scripts/antsRegistrationSyNQuick.sh). Coregistrated data were used to segment each brain using [SPM](https://www.fil.ion.ucl.ac.uk/spm/) and the GM, WM and CSF priors provided by Johnson *et al* to create the GM, WM and CSF probabilistic maps of each subject. Denoised and signal bias corrected images were used in parallel to create a study specific T_1w_ template, using [modelbuild](https://github.com/CoBrALab/optimized_antsMultivariateTemplateConstruction), an optimized pipeline using [antsMultivariateTemplateConstruction2](https://github.com/ANTsX/ANTs/blob/master/Scripts/antsMultivariateTemplateConstruction2.sh), an unbiased template building method developed in ANTs package. This method has been integrated to our local cluster ([ISLANDe Facilities](https://islande.hub.inrae.fr/)) and requires [qbatch](https://github.com/CoBrALab/qbatch) and [slurn](https://github.com/SchedMD/slurm). For computation, our cluster comprises 240 physical cores, each associated with 9.6GB of memory (2.3TB of memory in total), with 90TB of long-term storage and 72TB of high-performance storage. Once both linear and non-linear (flowfield maps) transformations were calculated for each animal, we compiled all the transformations calculated for each image and applied them once to denoised and signal bias corrected images using [antsApplyTransforms](https://antsx.github.io/ANTsRCore/reference/antsApplyTransforms.html) to limit interpolation effects. Resulting images have been used to create the TEBTA template by calculating the mean image of each normalized T_1w_ using [Ginkgo](https://framagit.org/cpoupon/gkg). Probabilistic maps (GM, WM, CSF) have been normalized using the linear and non-linear transformation provided by [modelbuild](https://github.com/CoBrALab/optimized_antsMultivariateTemplateConstruction) using the [antsApplyTransforms](https://antsx.github.io/ANTsRCore/reference/antsApplyTransforms.html) command and the TEBTA GM, WM and CSF priors have been created by calculating the mean image of each normalized map using [Ginkgo](https://framagit.org/cpoupon/gkg). Both templates and priors will be used for spatial normalization and segmentation for VBM analysis (see *Voxel Based Morphometry Analysis* section of main article).

##### *TEBTA atlas creation*

For the creation of the TEBTA atlas, the equine brain atlas provided by Johnson *et al* is used as a starting point. Firstly the Johnson’ template is linearly and nonlinearly coregistrated within the TEBTA space using [antsRegistrationSyNQuick](https://github.com/ANTsX/ANTs/blob/master/Scripts/antsRegistrationSyNQuick.sh). Then, both linear and non-linear transformations were applied to each ROI of the Johnson’ atlas using [antsApplyTransforms](https://antsx.github.io/ANTsRCore/reference/antsApplyTransforms.html). Each normalized ROI has been visually inspected to check boundaries and accuracy of registration. Then some regions of interest (ROIs) such like *Corpus Callosum*, *arbor vitae*, and ventricular systems have been updated/added using WM and CSF priors for a best fit to these ROIs to the TEBTA template. Eventually, additional subcortical ROIs such as septum, preoptic hypothalamic area, nucleus accumbens, striatum, etc. have been drawn and delimited manually using [fsleyes](https://fsl.fmrib.ox.ac.uk/fsl/fslwiki/FSLeyes) and [itksnap](http://www.itksnap.org/pmwiki/pmwiki.php) using both GM and WM priors maps to determine the boundaries of each new ROI.

To propose a valuable parcellation of the equine cortex, we implemented the methods previously used by Garin *et al* for the establishment of a functional atlas in lemur mouse^9^. Briefly, a multi-animal dictionary learning statistical analysis was performed with Nilearn (random_state = 0) on preprocessed rsfMR images (see *Functional Imaging Analysis* section of main article)^10^. A mask excluding the WM, CSF and subcortical areas was used to restrict the dictionary learning analysis to cortical functional data. During a pilot investigation, several analyses were performed using 30, 35, 40, 45, 50, 60, 90 and 120 sparse components. The study based on 60 sparse components was selected for the final analysis as it highlighted bilateral regions that matched well to anatomy or unilateral regions located to the sagittal midline (cingulate-related regions). Each bilateral component was split into two unilateral regions and labelled left or right. Regions smaller than 5 mm^3^ were excluded. This led to a 3D functional atlas composed by a mosaic of 55 local functional regions that were named using ITK-SNAP. The name of each ROI was defined using the names of brain structures reported by Schmidt *et al*^11^, the AAL2 human brain atlas but also using their structural connectivity.

##### *TEBTA fiber atlas creation: Structural connectivity of the equine brain*

Cortical ROIs resulting from the dictionary learning segmentation of the equine cortex were used to identify the largest bundle tracts of white matter to help for the identification of ROI based on their structural connectivity. For the construction of the fibre tractogram, we used the analytical Q-ball reconstruction model^12^ since this model relies on the decomposition of the diffusion-weighted signal into a modified spherical harmonics basis. We firstly computed fields of the orientation distribution functions (ODFs) stemming from the analytical Q-ball model including the fractional anisotropy map (FA), the mean diffusivity maps (MD), and the colour-encoded direction (CED) map. The analytical Q-ball model reconstruction was applied using spherical harmonics order 8 and a regularization factor λ = 0.006. Then, a streamline regularized deterministic (SRD) tractography algorithm^12^ was applied to the whole sample using the ODFs maps of aQBI previously computed, with the following parameters: 8 seeds per voxel, forward step 400 μm, maximum solid aperture angle 30°, minimum/maximum fibres length 10/300 mm. In the next step, tractogram were normalized to the TEBTA template using the previously calculated affine and diffeomorphic transformations (see *Diffusion Imaging Analysis* section of main article). Tractograms of each animal have been reduced by a random selection of 5% of total fibre count composing the tractogram and all the tractograms (n=23) were fused together and symmetrized to create a population-based tractogram. From this tractogram, we selected the large bundles (length>150mm) using a 2 steps approach. 1/ Bundles selection: in this step, bundles between each cortical ROIs have been selected by a ROI-to-ROI selection, labelled and merged. Then bundles between thalamus and each cortical ROI were selected as well as bundles between cerebellum and each cortical ROIs. With this selection, we expected to find long cortico-cortical pathways (i.e. cingulum), the thalamic projections such as the somatosensorial tract to identify the somatosensory areas and the cerebellocortical tract to identify the motor areas. To eliminate non-relevant fibres previously selected within the bundle tract we filtered them by length (>150mm) and by tortuosity (<2σ of mean of tortuosity). 2/ building of fibre atlas: here all the bundles selected have been visually inspected and 7 large bundle tracts (cingulum, corticospinal tract, anterior/posterior/inferior thalamic radiations, inferior longitudinal tracts and cerebellocortical tract) have been identified on basis of morphology, position within the brain, location and by comparison of fibre atlases available in humans and NHP. Areas connected to these tracts have been named according to their structural connectivity, position and literature^11^. All tractogram calculations and manipulation have been done by using [Ginkgo](https://framagit.org/cpoupon/gkg) toolbox developed by the Ginkgo team headed by C. Poupon. The Turone Equine Brain Template and Atlas resources are freely available on [Zenodo](https://zenodo.org/records/10731031).

### **Supplementary Results – Brain Imaging**

##### *Equine brain template creation and comparison.*

The TEBTA was built using the methods developed by the ANTs package. Comparison of signal to noise ratio (SNR) and contrast to noise ratio (CNR) between Johnson template and TEBTA are summarized in **Table S1**. SNR is higher for the TEBTA template (Jonhson=2.46 vs. TEBTA=8.03; +330%) while CNR is higher for the Johnson’ template (Jonhson=3.49 vs. TEBTA=1.72; -50%). Additionally, brain segmentation looks different between both resources. When the Johnson’ segmentation maps (FSL-FAST) seem to classify massively brain voxels in the white matter compartment (WM) rather than within grey matter compartment (GM), the TEBTA classification (Atropos) enlarged the cortical band and change voxels from WM to GM within caudate and mesencephalon (**Fig. S1a-b**, blue arrows). Additionally, the shape of ventricles looks different and are enlarged in TEBTA (**Fig. S1a-b**, orange arrows) probably because T_1w_ data have been acquired *in vivo* for TEBTA versus *ex vivo* in Johnson’ resources. Finally, some unexplained and extracephalic voxels classified as cerebrospinal fluid (CSF) are observed within the Johnson’ CSF prior but not in TEBTA CSF prior maps (**Fig. S1a-b**, red arrows).

##### *Equine brain atlas creation.*

A brain atlas is a cartography of the brain which is mandatory for the identification of brain territories involved in a biological or pathological process. The Johnson’ atlas is composed by a mosaic of 16 subcortical regions of interest (data not shown, see^8^) but no cortical regions have been segmented. To propose an accurate cortical parcellation of the equine brain, we used functional MRI data acquired at rest and use a multi-animal dictionary learning statistical approach to emulate the first functional cortical atlas of equidae. Using this approach, we identified 55 functional regions in the equine cortex. Among these regions, 9 were located on the sagittal midline and associated as frontal regions or cingulate cortex parts (anterior cingulate, anteromedial cingulate, cuneus/precuneus, frontal anteromedial, orbitofrontal, posterior cingulate, posteromedial cingulate, retrosplenial, superior frontal). The other regions were found bilaterally with an acceptable similar shape and location between both hemispheres. Naming of these functional regions was based on available literature20 and human AAL2. Then we conserved and normalised the 16 original subcortical labels from the Johnson atlas to the TEBTA. After visual inspection and manual correction, we added 48 subcortical regions (i.e. septum, preoptic hypothalamic area, nucleus accumbens, striatum, etc.) which have been drawn and delimited using GM and WM priors’ maps (**Fig. 2a** in the main article). Both cortical and subcortical labelling were finally merged to emulate a complete brain atlas composed by a mosaic of 119 ROIs (**Fig. 2b** in the main article). Using this atlas and tractograms calculated from the diffusion weighted imaging, we inferred the first fibre atlas of the equine brain which identifies the largest bundles tracts of white matter for the identification of the ROI based on their structural connectivity (**Fig. 2c** in the main article). We identified 7 large bundle tracts: the cingulum which permitted to identify cingulate cortices (anterior, medial, posterior) and to decipher frontal regions^6^, the corticospinal tract for the identification of the somatosensory regions^7^, the anterior thalamic radiations for the identification of the prefrontal region^5^, the posterior thalamic radiations for the identification of the occipital lobe^5^, the inferior thalamic radiations for the identification of the insula^5^, the inferior longitudinal tract one of the major occipitotemporal used for the identification of occipital areas^13^, and the cerebellocortical tract for the identification of motor regions^14^. Finally, ROI naming was terminated using neuroanatomical information found in literature and homologous unnamed regions were identified using ALL2 human brain atlas.

### **Supplementary information – Physiological Assessment**

#### **Assessment of endocrine and metabolic phycological markers**

##### *Plasma metabolites and hormones (insulin and IGF-1) assays*

The concentrations of plasma total cholesterol (Chol) and triglycerides (TGs) were measured by using the respective assays (References 80106 and 80019, Biolabo, Maizy, France). Plasma glucose levels were determined by using the GAGO-20 kit (Sigma Aldrich, Saint-Quentin Fallavier, France). Plasma insulin and IGF-1 were determined by using the equine insulin and IGF-1 enzyme-linked immunosorbent (ELISA) assays (MBS044785 and MBS017382; MyBioSource Inc, San Diego, CA) according to manufacturer's protocol. For all the measurements of these plasma metabolites and hormones, the inter-assay and intra-assay coefficients of variation were <8%.

##### *Oxytocin assay*

Oxytocin concentrations were determined by Enzyme ImmunoAssay (Enzo Life Sciences, ADI-901-153) after solid-phase extraction using Sep-Pack C18 cartridges (Waters, WAT054945). 1000µl of plasma samples were extracted and reconstituted in 230µL assay buffer. Measurements were performed in duplicate. The intra-assay coefficient of variation was 5.8 % at 40pg/mL and the assay sensitivity was 7.8pg/mL.

##### *Cortisol assays*

Cortisol concentrations were determined using competitive enzyme immunoassay after steroid extraction with a solvent (ethyl acetate/cyclohexane). Duplicate analyses were performed on each sample. The intra-assay coefficients of variation were 2.8% and 2.7% at 15ng/mL and 60ng/mL respectively. The assay sensitivity was 2ng/mL.

### **Supplementary Figures.**

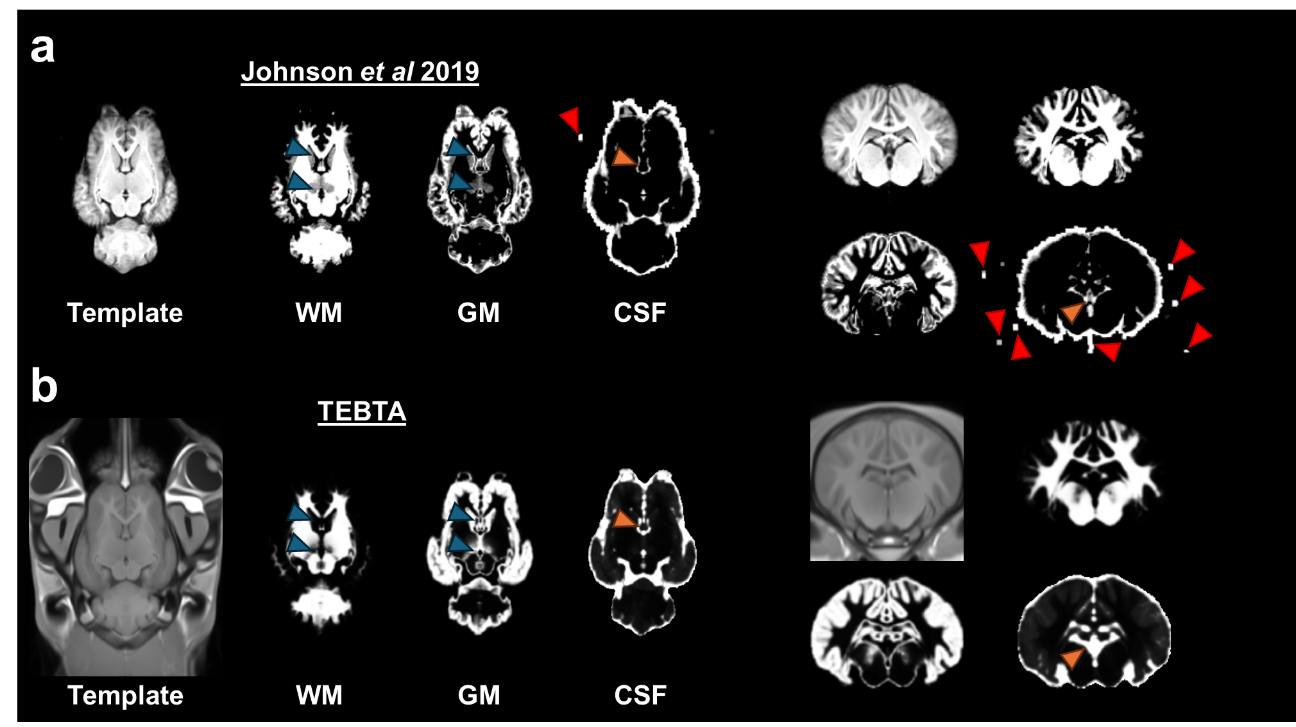

**Fig S1.** The Turone Equine Brain Template & Atlas. Sagittal and axial brain slices of the Johnson (**a**) and TEBTA (**b**) equine templates and their associated tissue probability maps of grey matter (GM), white matter (WM) and cerebrospinal fluid (CSF). All the images have been linearly co-registered within the same space for comparison. Blue arrows show the differences of classification between GM and WM through both resources. Oranges arrows show the differences of classification between GM/WM and CSF through both resources. Red arrows show extracephalic voxels misclassified as CSF.

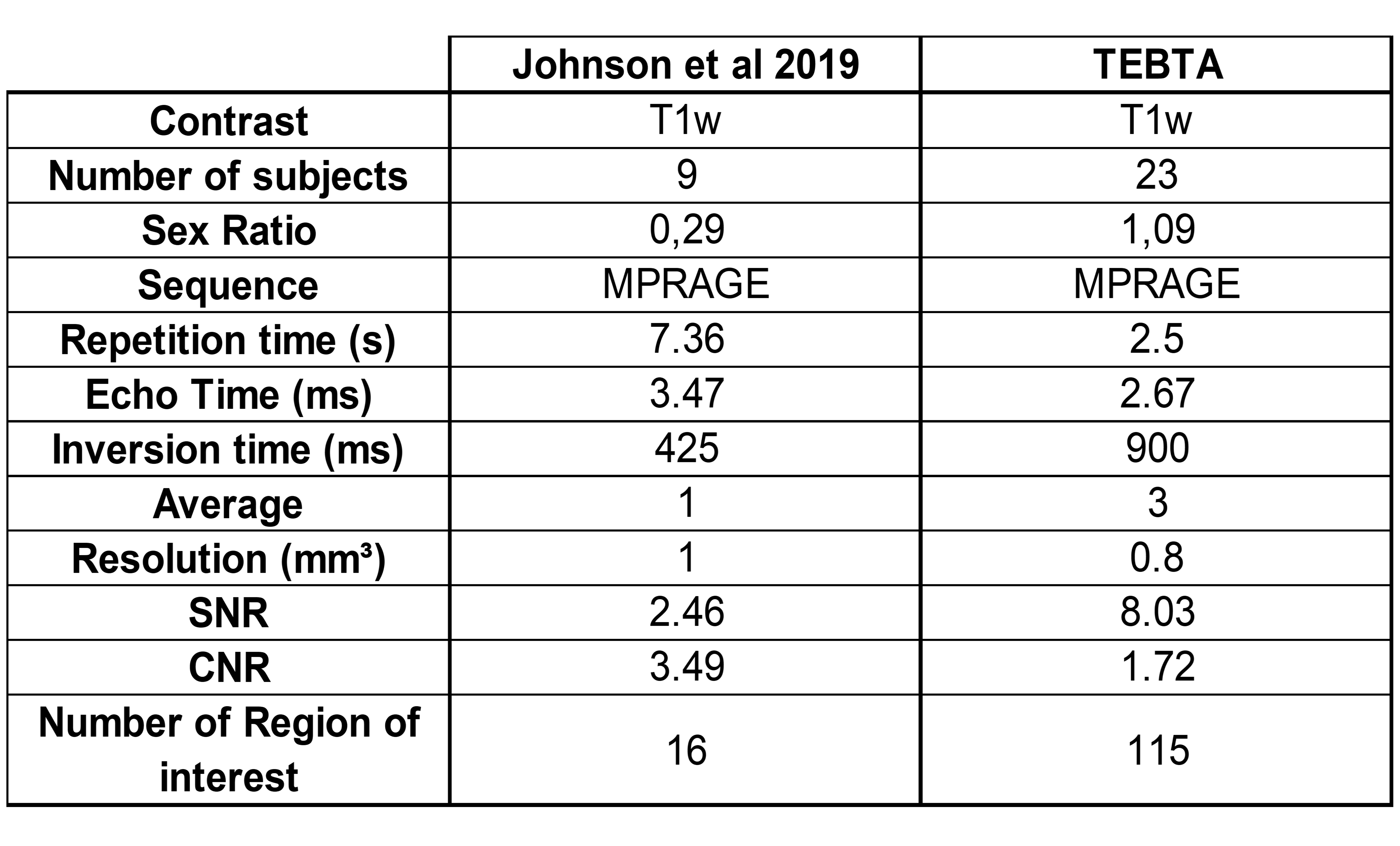

**Table S1.** Comparison of Johnson and Turone equine brain templates characteristics.

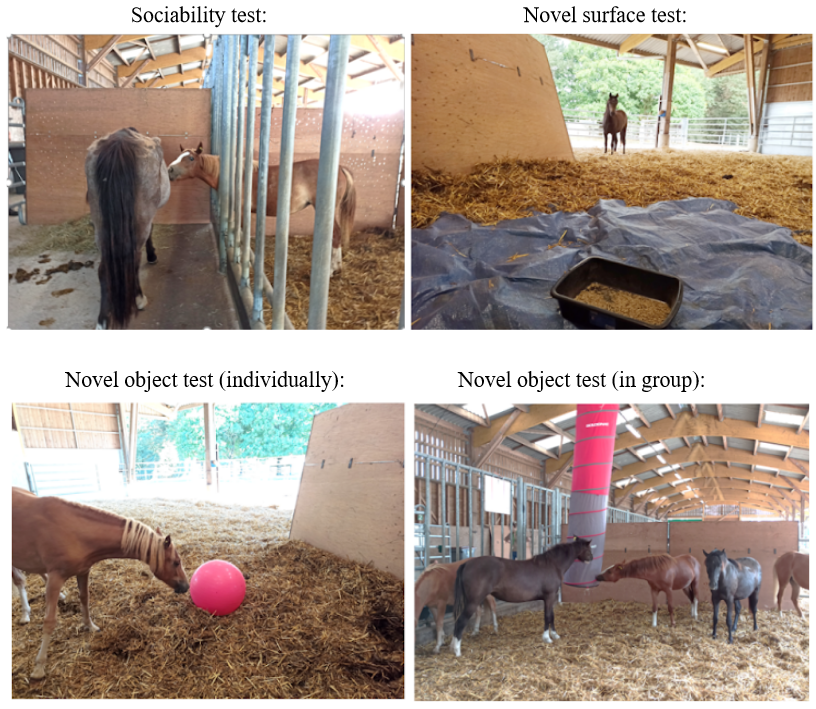

**Fig S2.** Pictures of the behavioural tests (Sociability test, Novel surface test, Novel object test in both individual or group situation).
